# Targeting the CDK4-Cyclin D Complex: A New Generation of Selective Kinase Inhibitors for Cancer Therapy

**DOI:** 10.64898/2026.09.18.752547

**Authors:** María Fuentes-Baile, José Antonio Encinar, Salomé Araujo-Abad, Elizabeth Perez-Valenciano, Pilar Garcia-Morales, Laura Fuertes-García, Camino de Juan Romero, Miguel Saceda

## Abstract

Glioblastoma (GBM) is an aggressive malignancy with limited therapeutic options and poor survival. Cyclin-dependent kinase 4 (CDK4) is overexpressed in GBM and promotes tumor progression through canonical cell-cycle regulation and extranuclear functions related to metabolism, invasion, and tumor microenvironment modulation. Current ATP-competitive CDK4/6 inhibitors show limited selectivity, potentially causing off-target effects and resistance. We developed a strategy to identify novel CDK4 inhibitors targeting the CDK4-Cyclin D1 protein-protein interaction, a key determinant of CDK4 activation. Virtual screening of approximately 950,000 compounds, combining molecular docking and molecular dynamics simulations, identified candidate molecules that were subsequently evaluated using proximity ligation assays, *in vitro* kinase activity assays, phosphokinase profiling, and functional analyses in patient-derived GBM cell lines and additional cancer models. Diophenic emerged as the lead candidate, demonstrating consistent antiproliferative activity across multiple GBM and epithelial cancer models. The compound selectively disrupted the CDK4-Cyclin D1 interaction while sparing the closely related CDK6 complex, and inhibited CDK4 kinase activity *in vitro*. Functional studies showed that diophenic induced substantial cell death in GBM cells, significantly reduced invasiveness, and exhibited distinct effects on cell migration compared with CDK4/6 inhibitors. Furthermore, phosphokinase profiling revealed that diophenic modulates a more restricted signaling network than ATP-competitive inhibitors, supporting greater functional selectivity and reduced off-target activity. Aiming the CDK4-Cyclin D1 interface represents a promising alternative to ATP-competitive CDK4/6 inhibition, which target a highly conserved site across the protein kinase superfamily. This approach provides a proof of concept for developing next-generation CDK4-selective therapeutics. Further studies should evaluate its translational potential in GBM.

## 1. Introduction

High-grade gliomas are aggressive brain tumors with poor prognosis, as glioblastoma (GBM) patients show a median survival of only 12 months [1]. Although Verhaak et al. (2010) classified GBMs into four molecular subtypes (proneural, neural, classical, and mesenchymal) [2], this molecular taxonomy has not yet translated into clinical practice, and therapeutic decisions still rely on biomarkers such as O6-methylguanine-DNA methyltransferase (MGMT) promoter methylation, isocitrate dehydrogenase 1/2 (IDH1/2) mutations, and 1p/19q codeletion [3].

In a previous study from our group on differential gene expression in GBM, one of the most consistent findings was the overexpression of Cyclin dependent kinase 4 (CDK4), a result also reported by other authors [4–6]. Based on this evidence, we hypothesized that CDK4 could represent a potential therapeutic target in GBM.

CDK4 plays a central role in cancer biology [7–9]. Its canonical function involves regulating the cell cycle through phosphorylation of the retinoblastoma (RB) protein, driving the G1-to-S transition. However, accumulating evidence has revealed several extranuclear functions of CDK4, including its participation in metabolic regulation, cell fate determination, cell dynamics, and modulation of the tumor microenvironment (TME) [10–12]. The CDK4-Cyclin D1 complex acts as a critical regulatory module that governs CDK4 nuclear-cytoplasmic localization and mediates its extranuclear activation [13,14].

Beyond the RB pathway, CDK4 promotes metabolic rewiring in cancer cells, favoring anabolic processes such as lipid biogenesis and glycolysis [15,16]. Pharmacological inhibition of CDK4/6 not only arrests the cell cycle but also induces senescence through the p16INK4A/RB pathway [17–22]. Moreover, the CDK4-Cyclin D axis contributes to tumor cell migration and invasion [9] and influences the TME by promoting a senescence-associated secretory phenotype (SASP) and enhancing T-cell infiltration [12,21]. Notably, CDK4 activity has also been linked to the regulation of PD-L1 expression and, based on this, they have an impact on the effectiveness of immunotherapy [23,24].

Despite the therapeutic relevance of CDK4/6 inhibition, these kinases share redundant functions, such as RB and FOXM1 phosphorylation [25]. The lack of selectivity of current CDK4/6 inhibitors complicates the interpretation of CDK4-specific effects [15,26,27]. Existing inhibitors (i.e. palbociclib, ribociclib, and abemaciclib) target the ATP-binding domain, a highly conserved motif among kinases, which results in off-target effects, toxicity, and the emergence of resistance mechanisms [15,25–27]. Furthermore, their combination with chemotherapeutic agents often leads to unpredictable, sequence-dependent outcomes. Thus, since they were designed to inhibit cell cycle entry, they would not be expected to cooperate with DNA-damaging or antimitotic agents. Although their collaboration with taxanes has been demonstrated in other types of cancer such as pancreatic adenocarcinoma (PAAD) [28,29]. Therefore, the sequence of administration of these inhibitors and other chemotherapeutic agents is important when evaluating their effects.

Given the conserved nature of the ATP-binding domain and the limited specificity of current inhibitors, the development of new compounds based on alternative targeting mechanisms is urgently needed. In this work, we propose a promising strategy for next-generation CDK4 inhibition which involves disrupting its interaction with Cyclin D1 (encoded by *CCND1*). This cyclin has been shown to be essential activating partner that determines CDK4 localization and function [13,14]. Targeting this protein-protein interface could provide enhanced specificity and functional selectivity by inhibiting CDK4’s oncogenic functions dependent on Cyclin D1, thereby minimizing off-target effects and improving the therapeutic index.

In this context, we performed a virtual screening of more than 900,000 compounds to identify small molecules capable of selectively disrupting the CDK4-Cyclin D1 interaction. Here, we report the discovery and characterization of diophenic, a novel protein-protein interaction (PPI) inhibitor that blocks CDK4 activity while sparing the closely related CDK6 complex. We demonstrate that diophenic exerts potent antiproliferative, pro-apoptotic, and anti-invasive effects across multiple cancer models, while modulating a substantially more restricted signaling network than conventional ATP-competitive inhibitors. Collectively, our findings validate the CDK4-Cyclin D1 interface as a therapeutically actionable target and support selective disruption of this interaction as a promising strategy for GBM and other CDK4-driven malignancies.

## 2. Methods

### 2.1. Chemical Libraries

The 3D structures of all computationally tested compounds were obtained from the MolPort database (https://www.molport.com/), which provides public access to compound structures in SMILES format. We developed in-house Python scripts to process and filter the initial compound library using the OpenBabel 3.1.1 and RDKit 2023 toolkits. The filtering criteria applied were: molecular weight ≤ 750 Da, ≤ 7 rotatable bonds, ≤ 4 chiral centers (only one stereoisomer), < 15 hydrogen bond acceptors, ≤ 5 hydrogen bond donors, topological polar surface area (TPSA) ≤ 175 Å², and no restriction on LogP values. While a LogP threshold of ≤ 3 is commonly used as a standard filter to improve drug-likeness, we omitted this constraint due to the hydrophobic nature of the target interface (CDK4-Cyclin D1 interaction). Anticipating potential limitations in docking performance with lower LogP compounds, we opted to include structures across a wider range of lipophilicities. After applying these filters, the final compound library comprised approximately 950,000 molecules.

### 2.2. Protein Structures

Several high-resolution structures of the human CDK4 enzyme (UniProt ID: P11802 - CDK4_HUMAN) are available, determined by X-ray crystallography and cryo-electron microscopy. These include complexes with various cyclins or with inhibitors bound to the ATP-binding site, such as abemaciclib in the 7SJ3 crystal structure. Upon comparison of structures 7SJ3 and 6P8E, we observed that the interface between CDK4-Cyclin D1 exhibits slight conformational changes in the presence of abemaciclib bound at the catalytic site. For this reason, we selected the 6P8E structure, which retains an unoccupied ATP-binding site, as the reference for our docking and dynamics studies. An additional rationale for selecting this structure is its practical applicability: both the recombinant CDK4-Cyclin D1 complex and the RB-derived peptide substrate are commercially available. This enables experimental validation of the computationally designed inhibitors through *in vitro* assays, as described below.

### 2.3. ADMET Prediction

Molecular descriptors -including TPSA, molecular weight (MW), the estimated logarithm (base 10) of solubility in molarity (cLogS), the estimated logarithm (base 10) of the partition coefficient between n-octanol and water (cLogP), the number of hydrogen bond acceptors (H-acceptors), hydrogen bond donors (H-donors), rotatable bonds, and the number of Lipinski’s Rule of Five violations (Ro5 violations)-were calculated using DataWarrior v5.0.0 software (Allschwil, Switzerland) [30,31]. The *in silico* prediction of absorption, distribution, metabolism, excretion, and toxicity (ADMET) properties were performed using the AdmetSAR web application [32] and DataWarrior v6.4.2 [33].

### 2.4. Molecular docking simulation

Molecular docking simulations were performed using YASARA Structure v25.1.13 (Vienna, Austria), employing the AutoDock 4 algorithm with the AMBER99 force field [30,31]. A cubic grid with dimensions of 23 × 23 × 23 Å³ was centered on the position occupied by co-crystallized Cyclin D1 on 6P8E, ensuring complete coverage of the CDK4 binding interface. The protonation states of CDK4 side chains at physiological pH (7.4) were assigned using the same software [34]. YASARA calculated the change in Gibbs free energy (ΔG, kcal/mol) for each docking pose, where more positive values indicate stronger binding. However, for consistency and ease of interpretation in this study, we report ΔG values as negative numbers, with lower values corresponding to stronger predicted binding affinities. The software output included molecular coordinates of the docked poses and their associated ΔG values. YASARA Structure was run on the high-performance Linux cluster hosted by the Scientific Computing Center at the Universidad Miguel Hernández (https://ccc.umh.es). All molecular graphics were prepared using PyMOL v3.0.1.

### 2.5. Molecular dynamics simulation

Molecular dynamics (MD) simulations were conducted using YASARA Structure v25.1.13 (Vienna, Austria) with the AMBER14 force field, as described in previous studies [31,35–37]. The simulation cell extended 20 Å beyond the protein in all directions and was solvated with water at a density of 0.997 g/mL. Initial energy minimization was performed under relaxed constraints using the steepest descent algorithm. Simulations were carried out under constant temperature (25 °C) and constant pressure conditions. To mimic a physiological environment, counterions (Na⁺ or Cl⁻) were added to neutralize the system, replacing water molecules to achieve a final NaCl concentration of 0.9%, and the pH was maintained at 7.4. Hydrogen atoms were added to ionizable groups based on calculated pKa values: a hydrogen atom was added if the computed pKa exceeded the simulation pH. pKa values were determined for each residue using the Ewald method [38]. All MD simulation steps were executed using the pre-installed *md_run.mcr* macro provided in the YASARA suite. Trajectories were saved every 100 ps. The molecular mechanics Poisson-Boltzmann surface area (MM/PBSA) method was used to estimate the alchemical binding free energy of drug candidates to CDK4, employing the *md_analyzebindenergy.mcr* macro, as previously described [31,35–37]. Compounds were retained only if they remained stably bound to CDK4 throughout the 100 ns simulation, with a root-mean-square deviation (RMSD) not exceeding 20 Å. Molecules exceeding this RMSD threshold were considered unstable and were excluded from further analysis. Additionally, compounds showing MM/PBSA binding energy values greater than −10 kcal/mol during the final 50 ns of the simulation were also discarded [39,40].

### 2.6. Cell culture

Our group at the Hospital General Universitario de Elche (HGUE, Elche, Spain) generated GBM cell lines derived from primary cultures, including HGUE-GB-16, -18, -37, -39, -40, -42, and -48 [41]. Hereafter, these cell lines are referred to as GB-followed by the corresponding cell line number. The exocrine PAAD cell lines IMIM-PC-2 and Hs766T, together with the colorectal carcinoma (COAD) cell line SW480, were provided by the Instituto Municipal de Investigaciones Médicas (IMIM, Barcelona, Spain). All cultures were maintained under conditions previously described [42]. Briefly, GBM cell lines were cultured in DMEM/F12 medium, while colon and pancreatic cell lines were cultured in DMEM, both media supplemented with 10% fetal bovine serum (FBS) and penicillin-streptomycin.

### 2.7. Proliferation assays

The antiproliferative activity of candidate CDK4 inhibitors was evaluated using the methylthiazolyldiphenyl-tetrazolium bromide (MTT) assay. Cells were seeded at 4000 cells per well in 96-well plates and pre-incubated for 24 hours at 37°C in a 5% CO_2_ atmosphere. Compounds were then added at increasing concentrations and incubated for 72 hours. Cell proliferation was determined by MTT assay, as described previously [42].

### 2.8. Immunofluorescence (IF)

A total of 35000 cells were seeded in 24-well plates on 12 mm glass coverslips. After 24 hours, cells were fixed with 4% paraformaldehyde (PFA) and blocked with FBS/phosphate buffered saline (PBS) (1x, 50 µL/mL). Primary antibodies against CDK4, CDK6 (1:100, mouse; NeoMarkers, CA, USA) and Cyclin D1 (1:100, rabbit; Cell Signaling, MA, USA) were applied. After washing, Alexa Fluor 568-conjugated anti-mouse and Alexa Fluor 488-conjugated anti-rabbit secondary antibodies (1:500; Invitrogen, Spain) were used. Nuclear staining was performed with DAPI. Coverslips were mounted in ProLong Gold Antifade Reagent (Invitrogen, Spain) and imaged using an LSM900 confocal microscope with Airyscan 2 (Carl Zeiss, Germany) at 63x magnification.

### 2.9. Proximity ligation assay (PLA)

For PLA, 35000 cells were seeded in 24-well plates on coverslips and treated with diophenic (10 µM, MolPort-001-025-593) for 6 hours. Cells were washed with PBS (1x), fixed with 4% PFA, washed again, permeabilized with 0.2% Triton X-100 in PBS (1x), and blocked with blocking solution for 1 hour at 37 °C before immunostaining. PLA was performed using the DuoLink *In Situ* kit (Merck, Spain) according to the manufacturer’s protocol. Primary antibodies against CDK4, CDK6 (1:100, mouse; NeoMarkers, CA, USA) and Cyclin D1 (1:100, rabbit; Cell Signaling, MA, USA) were applied. Samples were processed with DuoLink *In Situ* PLA Probe Anti-Mouse MINUS, DuoLink In Situ PLA Probe Anti-Rabbit PLUS, and DuoLink In Situ Red Detection Reagents (Merck, Spain). Red fluorescent spots corresponded to positive PLA signals, whereas nuclei were counterstained with DAPI. Negative controls were included by omitting one primary antibody. Images were acquired with an LSM900 confocal microscope with Airyscan 2 (Carl Zeiss, Germany) at 63x magnification, and quantification of red signals was performed with Fiji (ImageJ2).

### 2.10. CDK4 enzymatic activity

To assess CDK4 inhibition, a recombinant CDK4/Cyclin D1 complex (Life Technologies, CA, USA) and a peptide derived from RB protein (GenScript Biotech, Netherlands) were used. Kinase activity was measured with the ADP-Glo Kinase Assay (Promega, WI, USA), which quantifies ADP generated during the reaction, following the manufacturer’s instructions. Luminescence was recorded with a Cytation 3 microplate reader (BioTek, Germany).

### 2.11. Pathway reporter assay

Cells were seeded in 96-well plates at a density of 10000 cells/well in antibiotic-free medium. After 24 hours, cells were transfected with the pRb-TA-Luc, pE2F-TA-Luc, pMyc-TA-Luc, or pTA-Luc (negative control) reporter vectors from the Mercury™ Pathway Profiling Luciferase System 4 (Clontech, Mountain View, CA, USA). Transfections were performed using FuGENE® 6 Transfection Reagent (Promega, Madison, WI, USA) diluted in Opti-MEM™ I Reduced Serum Medium (Gibco, Life Technologies, Carlsbad, CA, USA) at a 3:1 reagent-to-DNA ratio. Each experimental condition was transfected in triplicate. 24 hours after transfection, cells were treated with diophenic (10 or 25 µM) and incubated for an additional 24 hours. Luciferase activity was measured using the ONE-Glo™ EX Luciferase Assay System (Promega, Madison, WI, USA). The reagent was added directly to the culture medium at a 1:1 (v/v) ratio, and luminescence was recorded after 3 minutes using a Cytation 3 Cell Imaging Multi-Mode Reader (BioTek, Germany).

### 2.12. Cell cycle analysis

Cells were cultured in 6-well plates and treated with 10 µM or 25 µM diophenic for 24, 48, or 72 hours. After treatment, cells were harvested by trypsinization and fixed in cold 75% ethanol at -20 °C for at least 1 hour. Fixed cells were pelleted and resuspended in 500 μL PBS containing 0.5% triton X-100, 25 μg/mL RNase A (Sigma Aldrich, MO, USA), and 25 μg/mL propidium iodide (PromoCell, Germany), then incubated for 30 minutes at room temperature in the dark. DNA content was analyzed with a BD FACSCanto™ flow cytometer (Becton Dickinson & Co., NJ, USA).

### 2.13. Cell death analysis

Cells were seeded in 6-well plates and treated under the following conditions: (i) 25 µM diophenic for 24, 48, or 72 hours; (ii) 10 µM diophenic, abemaciclib, or palbociclib for 48 hours; (iii) 25 µM diophenic with or without 25 µM Z-VAD-FMK (pan-caspase inhibitor, Calbiochem, Germany) for 24 hours. At the end of each treatment, cells were collected by trypsinization and incubated with 3 µM propidium iodide (PromoCell, Germany) for 30 minutes in the dark. Fluorescence was measured by flow cytometry with a BD FACSCanto™ flow cytometer (Becton Dickinson & Co., NJ, USA).

### 2.14. Gene silencing

For *CDK4* knockdown, cells were transfected with a specific siRNA targeting *CDK4*, using a non-specific (NS) siRNA as the negative control (Invitrogen, Carlsbad, CA, USA). Cells were seeded in 6-well plates in antibiotic-free culture medium and transfected with siRNA diluted in Opti-MEM™ I Reduced Serum Medium (Gibco, Life Technologies, Carlsbad, CA, USA). Lipofectamine™ RNAiMAX Transfection Reagent (Invitrogen, Carlsbad, CA, USA) was used at a final concentration of 2.5 µl/ml, together with 5 pmol/ml siRNA. Following transfection, cells were maintained for 72 hours at 37 °C in a humidified atmosphere containing 5% CO_2_. *CDK4* expression and cell cycle distribution were subsequently assessed.

### 2.15. Gene expression analysis

Total RNA was extracted using the NZY Total RNA Isolation kit (NZYtech, Lisbon, Portugal) following the manufacturer’s protocol. For cDNA synthesis, 1 µg of total RNA was reverse-transcribed using the High-Capacity cDNA Reverse Transcription Kit (Applied Biosystems, Foster City, CA, USA), according to the manufacturer’s instructions.

Quantitative real-time PCR (qPCR) was carried out in a final volume of 20 µl containing 4 µl of cDNA, 1 µl of the appropriate pre-designed TaqMan® Gene Expression Assay (Applied Biosystems, Foster City, CA, USA), and 10 µl of NZYSpeedy qPCR Probe Master Mix (2x), ROX plus (NZYtech, Lisbon, Portugal). The TaqMan® assays used were *CDK4* (Hs00364847_m1) and *GAPDH* (Hs02786624_g1), the latter serving as the endogenous control. Amplification was performed using a QuantStudio® 3 Real-Time PCR System (Applied Biosystems, Foster City, CA, USA).

### 2.16. Wound Healing Assay

Cell migration was evaluated by wound healing assays. Cells were seeded in 96-well plates and grown at 37 °C with 5% CO_2_ until confluence. A scratch was made with a 10 µl pipette tip, followed by PBS washing to remove detached cells. Cells were stained with Hoeschst 33342 and treated with 25 µM diophenic, abemaciclib, or palbociclib. Images were acquired at 0, 24, 48, and 72 hours using a Cytation 3 Cell Imaging Multi-Mode Reader (BioTek, Germany). Wound closure was quantified with ImageJ software.

### 2.17. Invasion Assay

Invasiveness was determined with the QCM™ 96-Well Cell Invasion Assay (Merck Millipore, MA, USA), based on the Boyden chamber principle. Cells were serum-starved for 24 hours, washed with PBS, trypsinized, and resuspended in media containing 5% bovine serum albumin (BSA). Suspensions of 1 x 10^6^ cells were incubated for 1 hour at 37 °C in the presence or absence of 25 µM diophenic or abemaciclib. Inserts with 8 µm pore polycarbonate membranes coated with ECMatrix™ were rehydrated with serum-free media 2 hours before the assay. Lower chambers were filled with either serum-free medium or medium containing 10% FBS, and cells were seeded in the upper chambers. After 24 hours, non-migrated cells were removed, and inserts were washed with PBS. Migrated cells were detached, incubated with lysis/staining buffer for 15 minutes at room temperature, transferred to a 96-well fluorescence plate, and quantified using a POLARstar Omega reader (BMG LabTech, Germany).

### 2.18. Phosphokinase array

Cells were treated with 25 µM diophenic or abemaciclib, or left untreated, for 3 hours. After treatment, cells were lysed and protein concentration was measured using the Pierce™ BCA Protein Assay Kit (Thermo Scientific™, MA, USA). Phosphorylation levels of 37 kinases were analyzed with the Proteome Profiler Human Phospho-kinase Assay Kit (R&D Systems, MN, USA), following the manufacturer’s protocol. Spot intensities were quantified using ImageJ software.

### 2.19. Statistical analysis

Descriptive analyses were performed with GraphPad Prism v9 (GraphPad Software, CA, USA). Mean and standard deviation (SD) values were calculated for all data. Normality was assessed using the Shapiro-Wilk test, and homoscedasticity with Levene’s test. Comparisons between two groups were carried out using Student’s t-test for parametric and homoscedastic data, whereas the Mann-Whitney U test was applied for non-parametric or heteroscedastic data. Differences were considered statistically significant at p < 0.05.

## 3. Results

### 3.1. Selection of CDK4 as a potential therapeutic target in glioblastoma

Previous transcriptomic analysis using microarray technology identified CDK4 as a potential target in GBM [4]. In order to confirm these results, we performed a RNA-seq transcriptomic analysis comparing RNA extracted from seven patients derived cell lines to healthy brain tissue. Accordingly, CDK4 was found to be increasingly overexpressed in primary GBM cultures (with a mixed population of tumor and non-tumoral cells) and GBM derived cell lines (**Fig. S1A**). To verify this observation, we analyzed CDK4 expression in GBM using publicly available datasets. Analysis of the GlioVis and UALCAN databases [43–45] showed a marked increase in CDK4 expression in GBM samples compared with non-tumoral brain tissue (**Fig. S1B,C**), suggesting that CDK4 inhibitors could represent a promising therapeutic strategy for this type of cancer.

Based on these results, CDK4 was selected for further investigation as a potential therapeutic target. Currently, three inhibitors of CDK4/6 approved by the European Medicines Agency (EMA) and the U.S. Food and Drug Administration (FDA) (i.e. palbociclib, ribociclib, and abemaciclib) are used primarily for hormone receptor-positive (HR+) and human epidermal growth factor receptor 2-negative (HER2−) breast cancer [46]. These ATP-competitive compounds block RB phosphorylation by targeting de CDK4/6 catalytic pocket, a site highly conserved across the kinase superfamily [47,48], which may contribute to off-target activity and clinical toxicity [49,50]. Since CDK4 activity requires Cyclin D1 binding, we reasoned that disrupting the CDK4-Cyclin D1 interface could offer a more selective therapeutic strategy than targeting the ATP-binding site [51] (see Graphical Abstract).

### 3.2. Computational approaches for the identification of novel CDK4 inhibitors

To identify potential CDK4 inhibitors targeting the Cyclin D1 interaction interface, a multi-step virtual screening strategy was applied to a chemical library of 954,861 commercially available compounds from MolPort (https://www.molport.com/). The selection process involved three sequential filters: 1) Molecular docking using the crystal structure of the human CDK4-Cyclin D1 complex (PDB ID: 6P8E) [52], yielding ΔG values from −11.3 to −8.9 kcal/mol (741 compounds retained). 2) ADMET prediction using DataWarrior and ADMETsar software (72 compounds passed). 3) MD simulations (100 ns) to select candidates with stable binding (maximum RMSD ≤ 20 Å) and favorable MM/PBSA binding energies (≤ −10 kcal/mol over the last 50 ns). This final step yielded 29 top candidates for further experimental evaluation (**Table S1** and **Fig. 1**), representing 0.003% of the initial library.

**Fig. 1.**
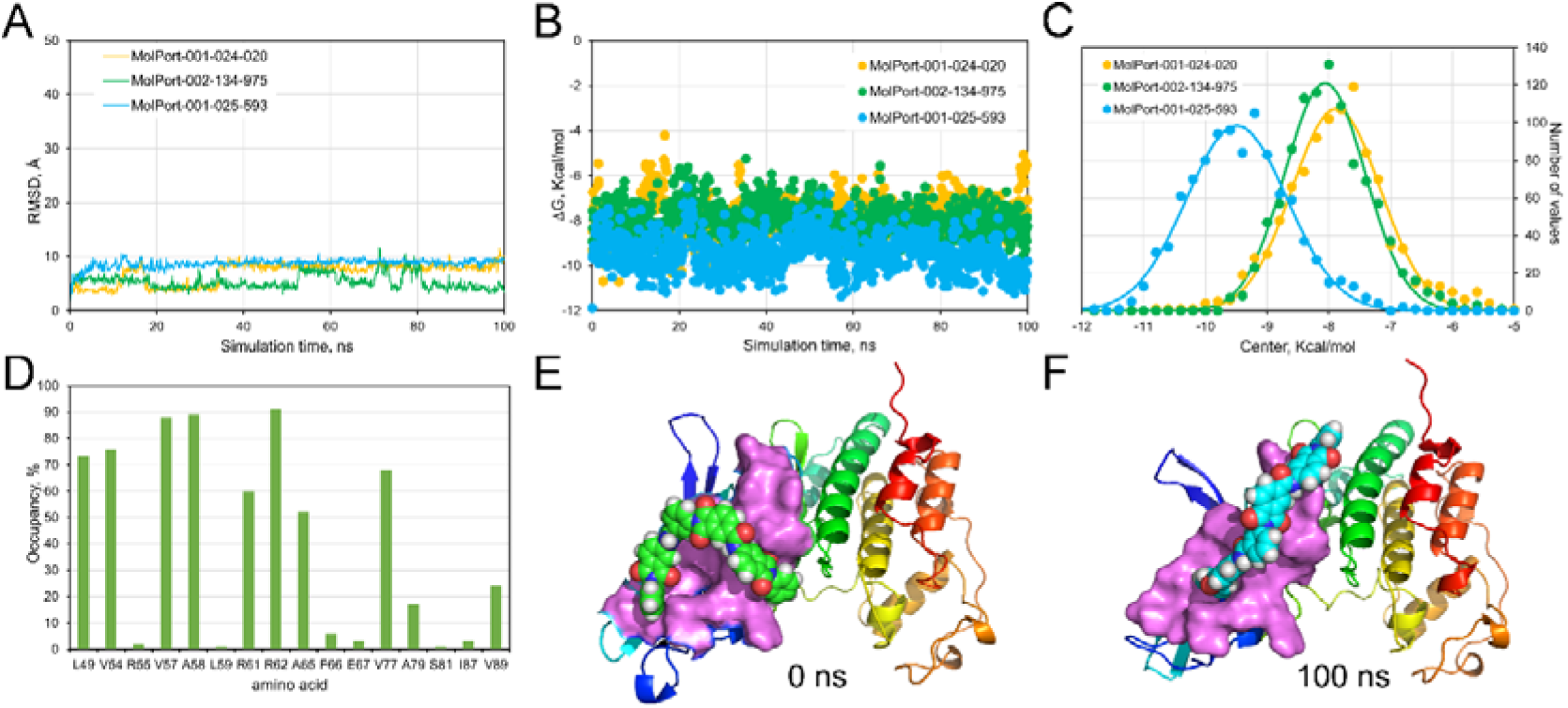
Computational screening of inhibitors targeting the CDK4-Cyclin D1 interaction. Molecular docking and dynamics simulations-based selection of compound diophenic (MolPort-001-025-593) as an inhibitor candidate of the CDK4-Cyclin D1 interaction. (**A**) trajectory of three analyzed compounds, with inhibitory effect of cell growth, bound to CDK4 throughout 100 ns MD simulation. (**B**) Autodock/vina calculated free energy variation (ΔG, Kcal/mol) for each 1000 snapshots of the same three compounds (Autodock/vina running under YASARA structure and calculated positively ΔG values indicate strong binding, however, here we flipped to negative value for simplicity). (**C**) frequency distribution of ΔG shown in (**B**) as a normal function and a continuous line for fitted Gauss equation. (**D**), Occupancy times of CDK4 residues involved in hydrogen bonding or hydrophobic interactions with diophenic during the 100 ns simulation. Representative binding poses of diophenic docked to the CDK4-Cyclyn D1 interface before (0 ns, (**E**) and after (100 ns, (**F**) the MD simulations are shown. The secondary structure of CDK4 is depicted in rainbow color from N-terminal (blue) to C-terminal (red). Surface residues interacting with diophenic are highlighted in magenta. H_2_O, Na⁺ and Cl⁻ ions were removed for clarity.

From these 29 candidates, we selected three structurally related molecules for biological validation based on their ability to inhibit GB-39 cell proliferation in a concentration-dependent manner (**Fig. 2**). These three compounds serve as representative examples of the selection criteria applied to all candidates in **Table S1** that successfully passed the three computational filters.

**Fig. 2.**
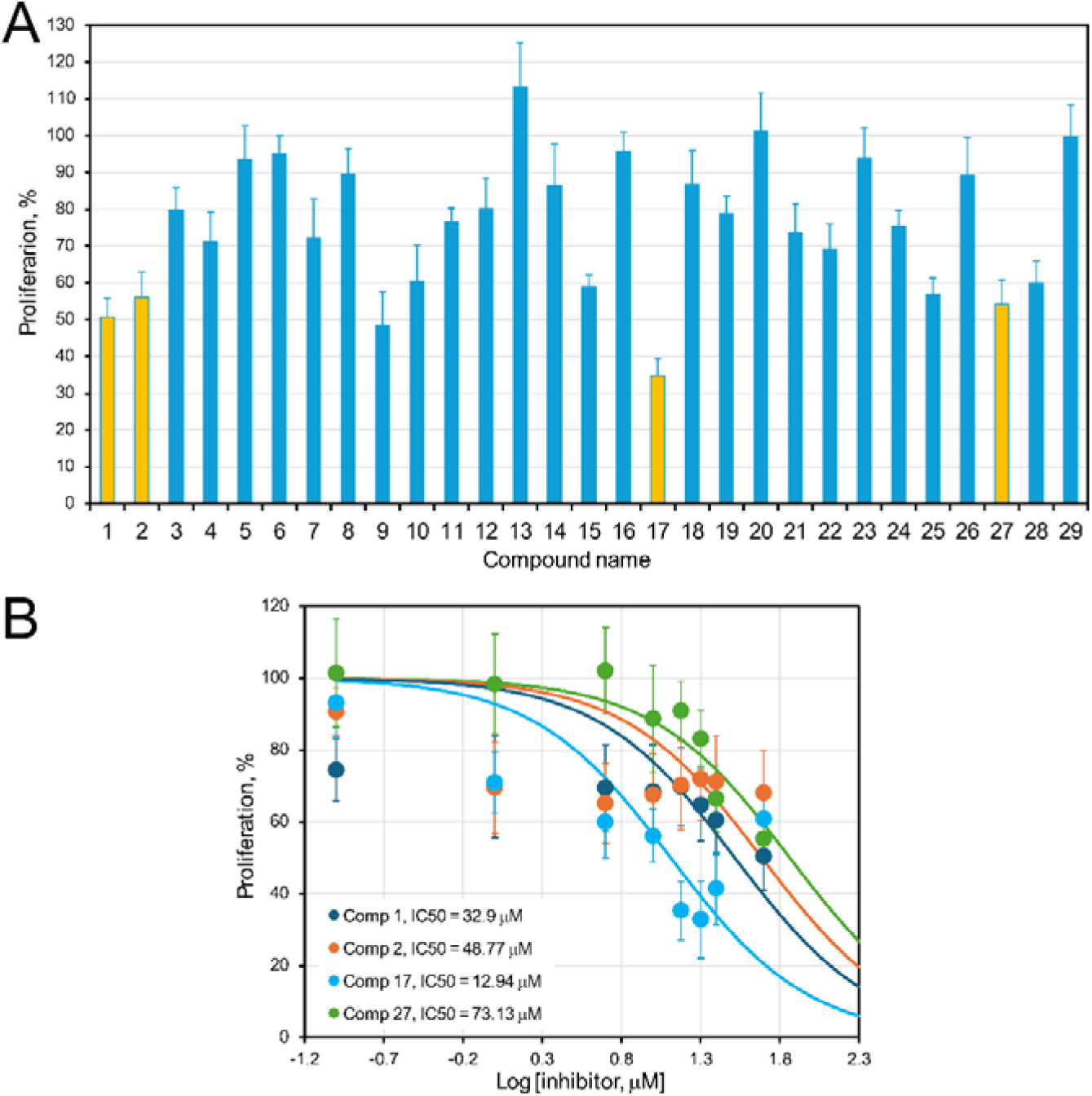
Effects of the best CDK4 inhibitor compounds on the proliferation of the GB-39 cell line. (**A**) Percentage of proliferation relative to the control after treatment with 25 µM of compounds 1-29. (**B**) The percentage of cell proliferation relative to untreated control is shown after treatment with compounds 1, 2, 17 (= diophenic), and 27 at concentrations ranging from 0.1 to 50 µM. Data are presented as mean ± SD (n = 6). Dose-response curves were fitted using a three-parameter logistic model [Y=100/(1+10^((X-LogIC50)), X: log of inhibitor concentration, and Y: normalized response, from 100 down to 0], and the legend indicates the corresponding IC_₅₀_ values.

While molecular docking provides a static model of ligand recognition at the protein binding site, MD simulations capture the time-dependent behavior of the molecular system, including ligand, receptor, water molecules, and ions, under physiological conditions. The MD simulation results were evaluated using the RMSD of each compound’s trajectory over 100 ns. As indicated in **Table S1** (fifth column), all selected candidates exhibited maximum RMSD values below 12 Å. Specifically, the three representative compounds demonstrated maximum RMSD values below 10 Å (**Fig. 1A**), suggesting stable binding conformations. However, considering that the CDK4-Cyclin D1 interaction surface exceeds 20 Å², RMSD values between 12 and 15 Å, as observed for other candidates in **Table S1**, may still represent valid binding events.

MM/PBSA binding free energy values calculated with YASARA were used to estimate binding affinity. The sign of these values was inverted so that more negative values correspond to stronger binding. To complement this analysis, we computed ΔG values (kcal/mol) using Vina for snapshots extracted from the MD simulations (**Fig. 1B**). These values were analyzed through frequency distributions fitted to a Gaussian function: Y=Amplitude*exp(-0.5*((X-Mean)/SD)^2^), showing more negative ΔG values which indicate stronger ligand-receptor interactions (**Table S1**, **Fig. 1C**.).

Finally, we evaluated for each compound the frequency of interactions between its functional groups and CDK4 residues throughout the simulation obtaining their interaction “fingerprint”. When analyzing the percentage occupancy of CDK4 residues interacting with diophenic (MolPort-001-025-593) (**Fig. 1D**), we observed that residues L49, V54, V57, A58, A65, V77, and V89 maintained hydrophobic interactions with diophenic for more than 20% of the simulation time (**Fig. 1E,F**). After an initial equilibration phase (first 5 ns, **Fig. 1A**), diophenic reorients within the binding site and adopts the conformation depicted in **Figure 1F**, engaging with hydrophobic residues on the CDK4 surface (shown in violet; ligand shown as van der Waals spheres).

### 3.3. Biological screening of candidate compounds on GBM

To evaluate the effect of the 29 candidate compounds on cellular models, MTT assays were first performed on the GBM cell line GB-39. Increasing concentrations of all compounds were tested, showing a highly variable response among the different compounds as can be seen by their effects on cell proliferation at 25 µM (**Fig. 2A**). The compounds 1, 2, 17, and 27 (highlighted in light orange) were showing the most pronounced inhibitory effects and therefore were selected to further investigate their dose-response using a broader concentration range (**Fig. 2B**). From these assays, IC_50_ values were determined to be 32.9 µM, 48.77 µM, 12.94 µM, and 73.13 µM, respectively. Since compound 17 exhibited the strongest antiproliferative activity, we designated it as diophenic for further reference and focused on it for the subsequent experiments in this study.

Next, we extended the analysis of diophenic to the remaining GBM patient-derived cell lines available in our laboratory (GB-16, GB-18, GB-37, GB-40, GB-42, and GB-48) (**Fig. 3A**), as well as to other cancer cell models, including IMIM-PC-2 and Hs766T (PAAD) and SW480 (COAD) (**Fig. 3B**). In all cases, a consistent, concentration-dependent decrease in cell proliferation was observed. Dose-response curves for the GBM cell lines were plotted on a semi-logarithmic scale, and IC_50_ values were calculated for each cell line (**Fig. S2**). The IC_50_ values obtained in this expanded analysis were consistently lower than those initially determined for diophenic, with the lowest value observed in GB-42 (0.96 ± 0.05 µM). Overall, these findings indicate that diophenic exerts a broad antiproliferative effect across GBM and epithelial cancer cell models.

**Fig. 3.**
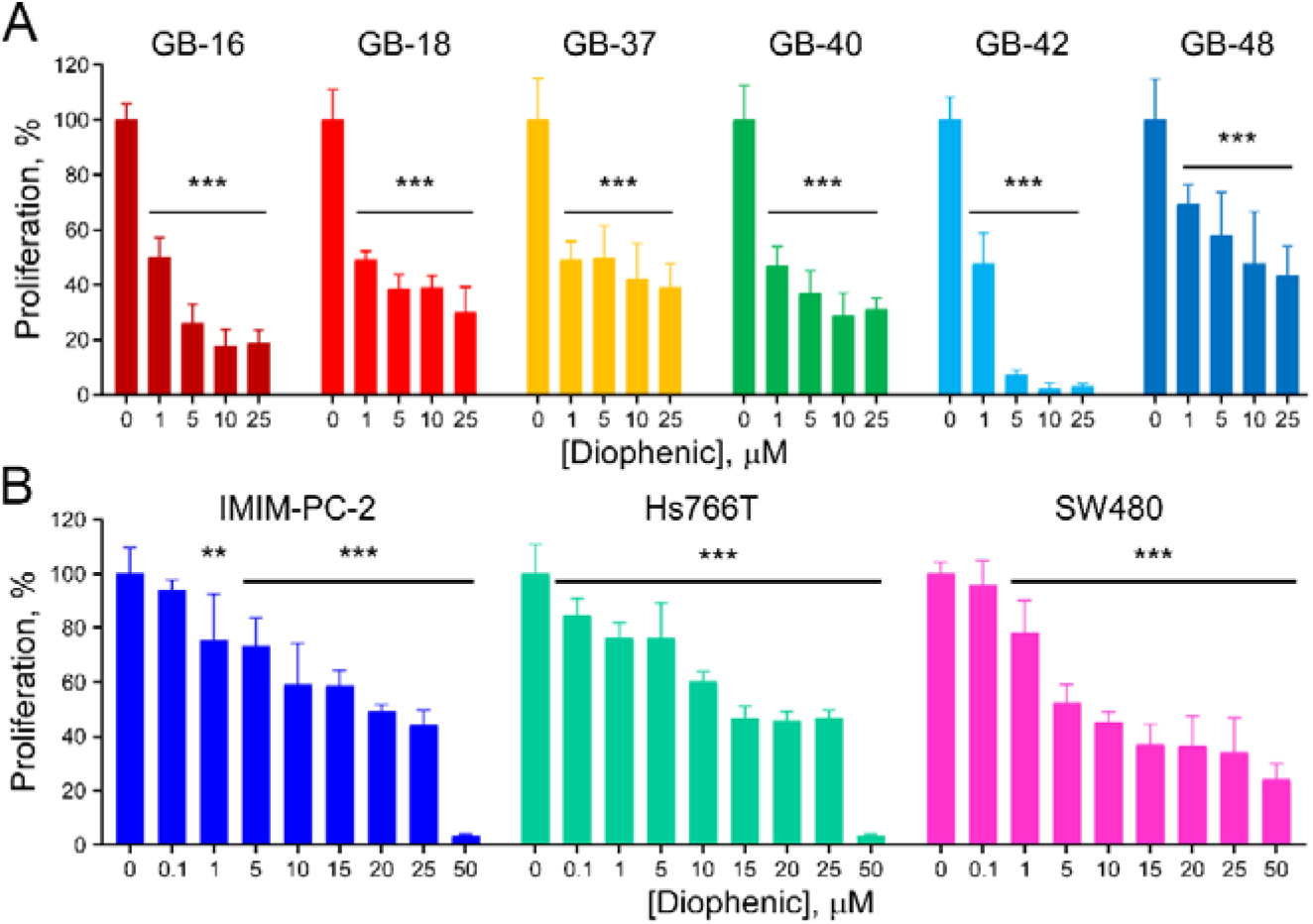
Effects of CDK4 inhibitor diophenic on cell proliferation. (**A**) Percentage of proliferation relative to the control after treatment with 1 to 25 µM diophenic in GBM cell lines. (**B**) Percentage of proliferation relative to the control after treatment with diophenic at concentrations ranging from 0.1 to 50 µM in IMIM-PC-2, Hs766T, and SW480 cell lines. Results are shown as mean ± SD, with n=6. ** *p* ≤ 0.01, *** *p* ≤ 0.001.

### 3.4. Diophenic blocks the interaction between CDK4 and Cyclin D1 in cellulo

To determine whether the interaction between endogenous CDK4 and Cyclin D1 occurs within cancer cells, we analyzed several tumor cell lines. For GBM, we used the GB-39 and GB-42 cell lines, and for COAD, we included the SW480 cell line.

We first performed Immunofluorescence (IF) experiments to examine whether both proteins were expressed and colocalized in the same cellular compartments (**Fig. S3**). CDK4 was highly expressed in the nuclei of GBM cells, as indicated by its colocalization with DAPI, but was also present in the cytoplasm. In contrast, CDK4 was predominantly cytoplasmic in COAD cells. Similarly, Cyclin D1 was expressed in both the cytoplasm and the nucleus of all tested cell lines. Quantification of colocalization using the Pearson’s correlation coefficient confirmed a higher level of co-expression in GBM cells.

We next assessed the *in cellulo* interaction between the two proteins using the *in situ* Duolink proximity ligation assay (PLA), which detects PPIs occurring at distances below 16 Å (**Fig. 4A**). Red fluorescent puncta corresponding to PLA signals indicated that CDK4 efficiently interacted with Cyclin D1, mainly within the nucleus, in all untreated control cells, consistently with the IF data. To confirm that this interaction was disrupted by the inhibitory compound diophenic, all cell lines were treated with 10 µM diophenic for 6 hours.

**Fig. 4.**
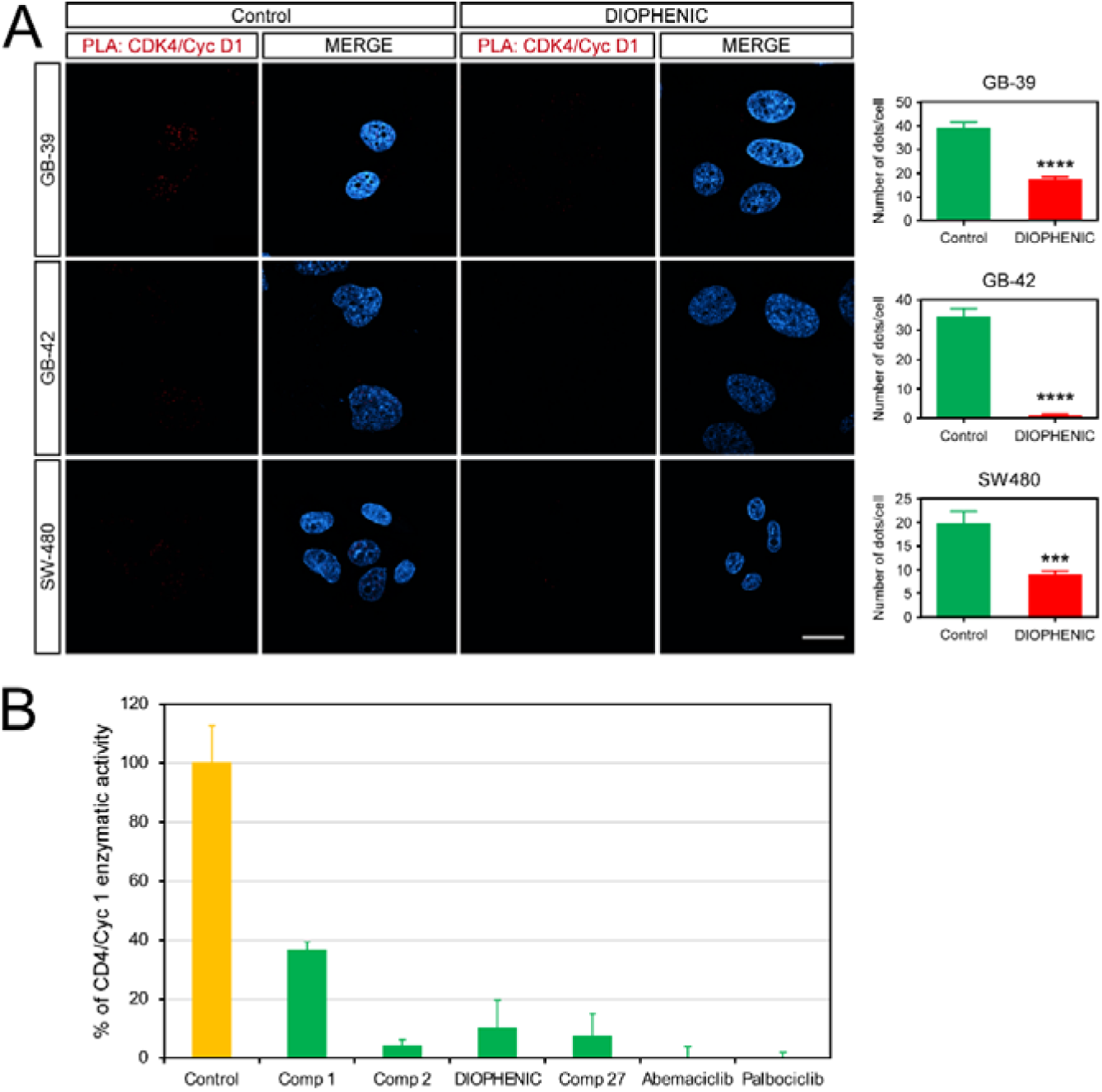
Effect of diophenic on the interaction between CDK4 and Cyclin D1. (**A**). Proximity ligation assay (PLA) performed in GB-39, GB-42, and SW480 tumor cells treated with 10 µM diophenic for 6 h. The scale bar represents 10 µm. (**B**). Percentage of enzymatic activity resulting from CDK4-CYCLIN D1 interaction after treatment with 50 µM of C1, C2, diophenic, C27, abemaciclib, or palbociclib for 1 h. Data are presented as mean ± SD (n = 3). *** *p* ≤ 0.001

In the GB-42 line, diophenic was particularly effective, nearly abolishing the nuclear CDK4-Cyclin D1 interactions. In contrast, the effect was less pronounced in GB-39. A marked reduction in CDK4-Cyclin D1 interaction was also observed in SW480 cells, suggesting that diophenic may exert inhibitory activity not only in GBM but also in other tumor types.

To evaluate the specificity of diophenic for CDK4 over its closely related homolog CDK6, we examined the CDK6-Cyclin D1 interaction in GB-42 cells by IF and PLA (**Fig. S4**). CDK6 and Cyclin D1 showed expression and colocalization patterns comparable to those observed for CDK4 (**Fig. S4A**). However, in contrast to the CDK4-Cyclin D1 interaction, PLA signals corresponding to the CDK6-Cyclin D1 interaction remained unchanged following treatment with diophenic (**Fig. S4B**). These results indicate that diophenic selectively disrupts the CDK4-Cyclin D1 interaction without affecting the closely related CDK6-Cyclin D1 complex.

To determine whether disruption of the CDK4-Cyclin D1 interaction translates into functional changes in the downstream RB pathway, GB-42 cells were transiently transfected with Rb-, E2F-, or Myc-responsive luciferase reporter constructs, or with the promoterless pTA-Luc vector as a negative control, and subsequently treated with diophenic (10 or 25 µM) for 24 hours (**Fig. S5**). Rb reporter activity was significantly reduced at both concentrations, by approximately 30% and 55% at 10 and 25 µM, respectively, indicating an increase in RB-mediated transcriptional repression. This finding is consistent with enhanced RB activity resulting from reduced CDK4-mediated RB phosphorylation. E2F reporter activity was significantly reduced only at 25 µM, by approximately 35%, suggesting that a higher concentration of diophenic is required to produce a measurable reduction in E2F transcriptional activity. Myc reporter activity was also significantly reduced at both concentrations, consistent with decreased E2F-dependent transcription of MYC downstream of RB reactivation. Overall, these results indicate that disruption of the CDK4-Cyclin D1 interaction by diophenic is accompanied by enhanced RB-mediated transcriptional repression and reduced E2F- and Myc-dependent transcriptional activity, consistent with inhibition of the RB-E2F-Myc regulatory axis.

Finally, to determine whether diophenic and the other selected compounds directly affect CDK4 kinase activity, we performed an *in vitro* kinase activity assay using the recombinant CDK4/Cyclin D1 complex and an RB protein peptide as substrate. Both the candidate and commercial inhibitors were tested at 50 µM for 1 hour, and ADP production was used as a redout of kinase activity. The results showed that the tested compounds displayed inhibitory capacities comparable to those of the commercial inhibitors abemaciclib and palbociclib, with compound 1 showing the weakest effect, reducing CDK4 activity by approximately 60% (**Fig. 4B**). These findings provide complementary biochemical evidence that the selected compounds inhibit CDK4 kinase activity.

Together, these results confirm that diophenic selectively disrupts the CDK4-Cyclin D1 interaction without affecting the closely related CDK6-Cyclin D1 complex. This effect is accompanied by enhanced RB-mediated transcriptional repression and reduced E2F- and Myc-dependent transcriptional activity, consistent with inhibition of the CDK4-RB-E2F regulatory axis. Moreover, the *in vitro* kinase assay confirms that diophenic and the other selected compounds can inhibit CDK4 kinase activity, further supporting their potential as CDK4-targeting compounds.

### 3.5. Diophenic induces cell death in GBM cell lines

To determine whether the reduction in cell proliferation previously observed with diophenic (**Fig. 2A, 3A**) was due to a cytostatic or cytotoxic effect, we analyzed cell-cycle distribution in GB-39 and GB-42 cells treated with 10 µM or 25 µM diophenic for 24, 48, and 72 hours (**Fig. 5A, B**). In both cases, the most striking effect was an increase in the percentage of cells in the subG1 phase, which is associated with DNA fragmentation and cell death. At 10 µM, a gradual accumulation of cells in subG1 was observed over time in both cell lines while at the higher concentration, that population appeared to decrease at longer exposure times.

**Fig. 5.**
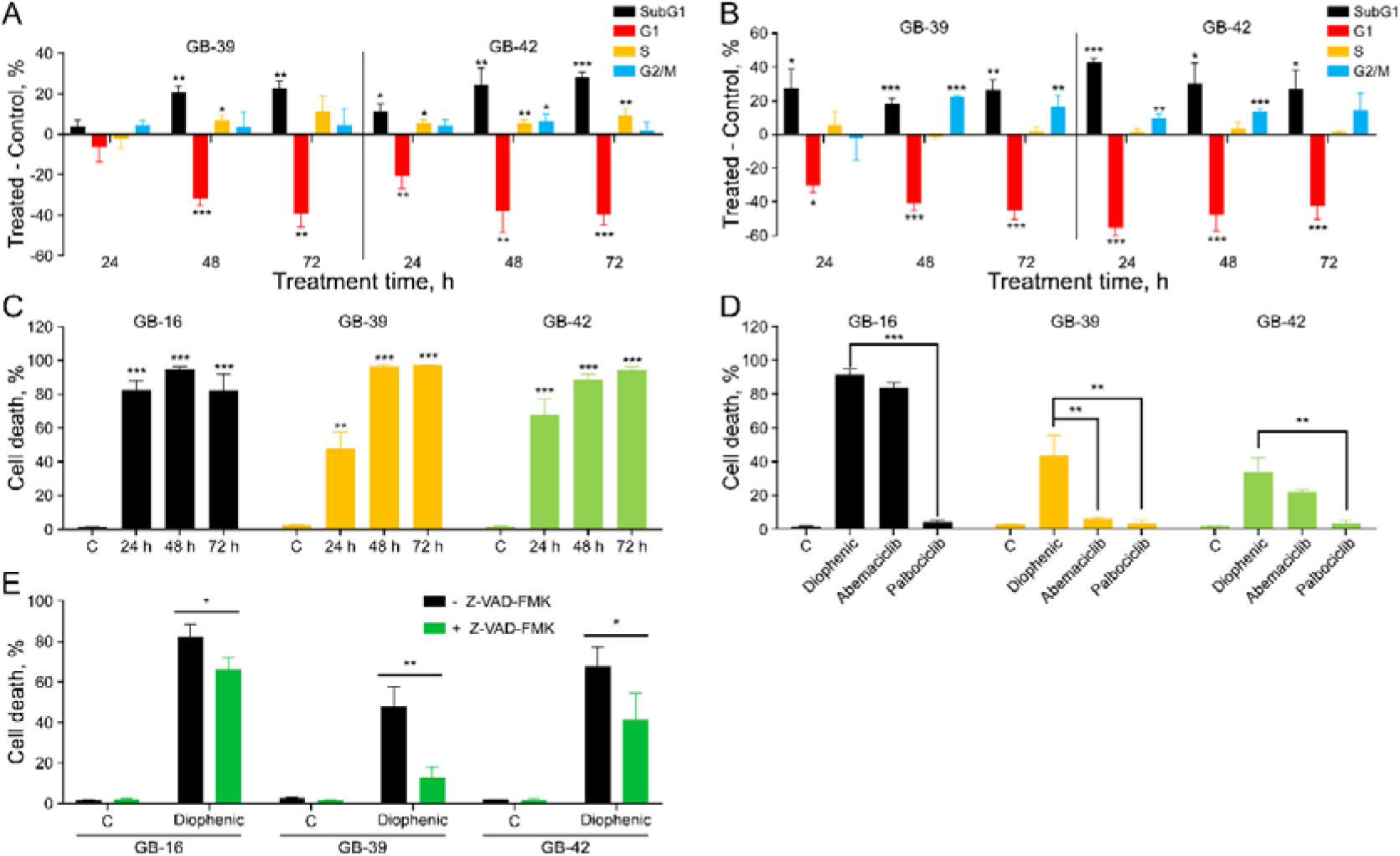
Diophenic-induced cell death in GBM cell lines. GB-39 and GB-42 cells were treated with 10 µM (**A**) or 25 µM (**B**) diophenic for 24, 48, and 72 h. Cell cycle distribution was analyzed by flow cytometry. The graphs show the difference in the percentage of cells in each phase relative to the untreated control. GB-16, GB-39, and GB-42 cells were treated with 25 µM diophenic for 24, 48, and 72 h (**C**), 10 µM diophenic, abemaciclib, or palbociclib for 48 h **(D**); and 25 µM diophenic in the presence or absence of 25 µM of a pan-caspase inhibitor (Z-VAD-FMK) for 24 h (**E**). Cell death percentage was assessed by flow cytometry using propidium iodide staining). Data are presented as mean ± SD (n = 3). * *p* ≤ 0.05, ** *p* ≤ 0.01, *** *p* ≤ 0.001.

To further investigate whether the cell-cycle alterations induced by diophenic were related to CDK4 inhibition, we performed *CDK4* silencing using siRNA in GB-16, GB-39, and GB-42 cells (**Fig. S6**). *CDK4* mRNA expression was reduced by more than 97% in all three cell lines compared with the non-specific (NS) control, confirming an efficient and consistent knockdown (**Fig. S6A**). Analysis of cell-cycle distribution following *CDK4* silencing revealed a cell-line-dependent phenotype. GB-16 showed an increase in the S and G2/M populations, whereas GB-39 cells exhibited a predominantly cytotoxic phenotype, characterized by a marked increase in the subG1 population. In contrast, GB-42 cells showed a clear accumulation of cells in G1, consistent with the canonical role of CDK4 in promoting the G1/S transition (**Fig. S6B**). Notably, the increase in the subG1 population observed following *CDK4* silencing in GB-39 is consistent with the cytotoxic response induced by diophenic, which also increased the subG1 population (**Fig. 5A, B**). The heterogeneous responses observed across cell lines, particularly between genetic and pharmacological CDK4 inhibition, may reflect differences in the kinetics, magnitude, and duration of CDK4 inhibition, as well as cell-line-specific mechanisms of adaptation.

Because cells with severely fragmented DNA may fall outside the detectable population in cell-cycle analyses, we repeated the experiment using an alternative method to assess plasma membrane disruption as a marker of cell death. In this case, GB-16 (the most sensitive line in the MTT assays), GB-39, and GB-42 cells were treated with 25 µM diophenic for 24, 48, and 72 hours. In GB-16 cells, approximately 80% cell death was detected after just 24 hours of treatment. In the other two lines, cell death increased progressively over time, reaching nearly 100% after 72 hours (**Fig. 5C**). These results confirm that diophenic induces cell death in GBM cells, with GB-16 being the most sensitive line to treatment.

Next, we compared the cytotoxic effect of diophenic with that of the commercial CDK4/6 inhibitors abemaciclib and palbociclib. GB-16, GB-39, and GB-42 cells were treated with 10 µM of each compound for 48 hours, and plasma membrane integrity was analyzed (**Fig. 5D**). The results were cell context dependent. In GB-16 cells, diophenic induced approximately 90% cell death, compared with ∼85% for abemaciclib and only 4% for palbociclib. In GB-39 cells, diophenic caused around 45% cell death, significantly higher than the <10% observed with the commercial inhibitors. Finally, in GB-42 cells, diophenic induced ∼35% cell death, abemaciclib ∼20%, and palbociclib ∼3.5%. In all cases, diophenic was the compound that induced the highest level of GBM cell death.

To investigate whether diophenic-induced cell death was caspase-dependent, cells were treated with the pan-caspase inhibitor Z-VAD-FMK (**Fig. 5E**). In all three GBM cell lines tested, Z-VAD-FMK significantly reduced diophenic-induced cell death, with the most marked reduction (∼75%) observed in GB-39 cells. These findings indicate that diophenic triggers, at least in part, a caspase dependent apoptotic cell death mechanism in GBM cells.

### 3.6. Diophenic reduces cell invasiveness and exhibits greater specificity than abemaciclib

To further explore the antitumor effects of diophenic, we analyzed its impact on cell migration and invasiveness. We first performed wound-healing assays in GB-39, GB-42, and SW480 cells treated with 25 µM diophenic, abemaciclib, or palbociclib (**Fig. 6A, Fig. S7**). Both commercial CDK4/6 inhibitors, abemaciclib and palbociclib, significantly reduced migration in all three cell lines, while diophenic did not alter cell migration under the same conditions.

**Fig. 6.**
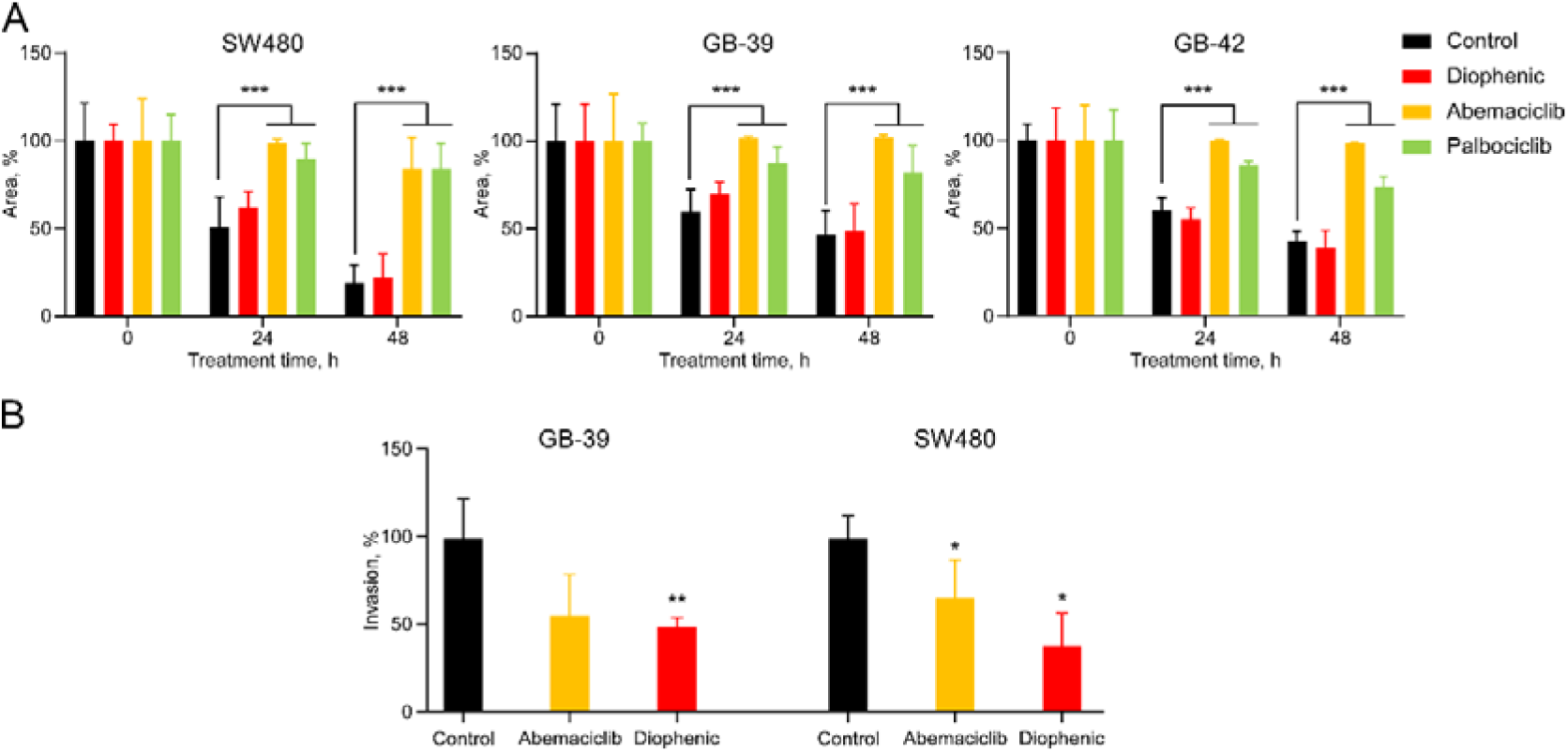
Effect of CDK4 inhibitors on cell migration and invasiveness. (**A**) GB-39, GB-42, and SW480 cells were treated with 25 µM diophenic, abemaciclib, or palbociclib for 48 h and labeled with Hoechst. Images were taken at baseline, 24, and 48 h, and wound area was quantified using ImageJ. (**B**) GB-39 and SW480 were treated with 10 µM diophenic or abemaciclib for 24 h, and cell invasiveness was assessed. Data are presented as mean ± SD of the percentage of invasion relative to control (n = 6). * *p* ≤ 0.05, ** *p* ≤ 0.01, *** *p* ≤ 0.001.

Next, we evaluated cell invasiveness in GB-39 and SW480 cells treated with 10 µM diophenic or abemaciclib. Abemaciclib caused a 35-45% reduction in invasiveness, whereas diophenic led to a stronger decrease, ranging from 50-60% (**Fig. 6B**). These results indicate that, unlike the commercial CDK4/6 inhibitors, diophenic affects cell invasiveness, without altering migration.

As previously mentioned, the commercial inhibitors target the conserved ATP-binding domain of CDK4, whereas diophenic interferes with the CDK4-Cyclin D1 interaction. Therefore, to determine whether or not this differential effect between abemaciclib, palbociclib, and diophenic could be related to their distinct mechanisms of action, we next performed a phosphokinase array to identify which signaling pathways were affected by each compound.

To compare the signaling effects of diophenic and abemaciclib, we performed a phosphokinase array using GB-42 cells treated with 25 µM of each compound for 3 hours (**Fig. 7**). We observed that several kinases were affected exclusively by abemaciclib, including MSK1/2, PLC-γ1, Src, Yes, P70S6K, and RSK1/2/3. Moreover, a second group of kinases was modulated by both compounds, but to a greater extent by abemaciclib, such as eNOS, Lyn, Akt, c-Jun, PYK2, STAT3, and HSP60. Interestingly, although both inhibitors similarly affected STAT5a/b, β-catenin, and p53 (Ser15), only diophenic modified the phosphorylation of GSK-3α/β and p53 (Ser392).

**Fig. 7.**
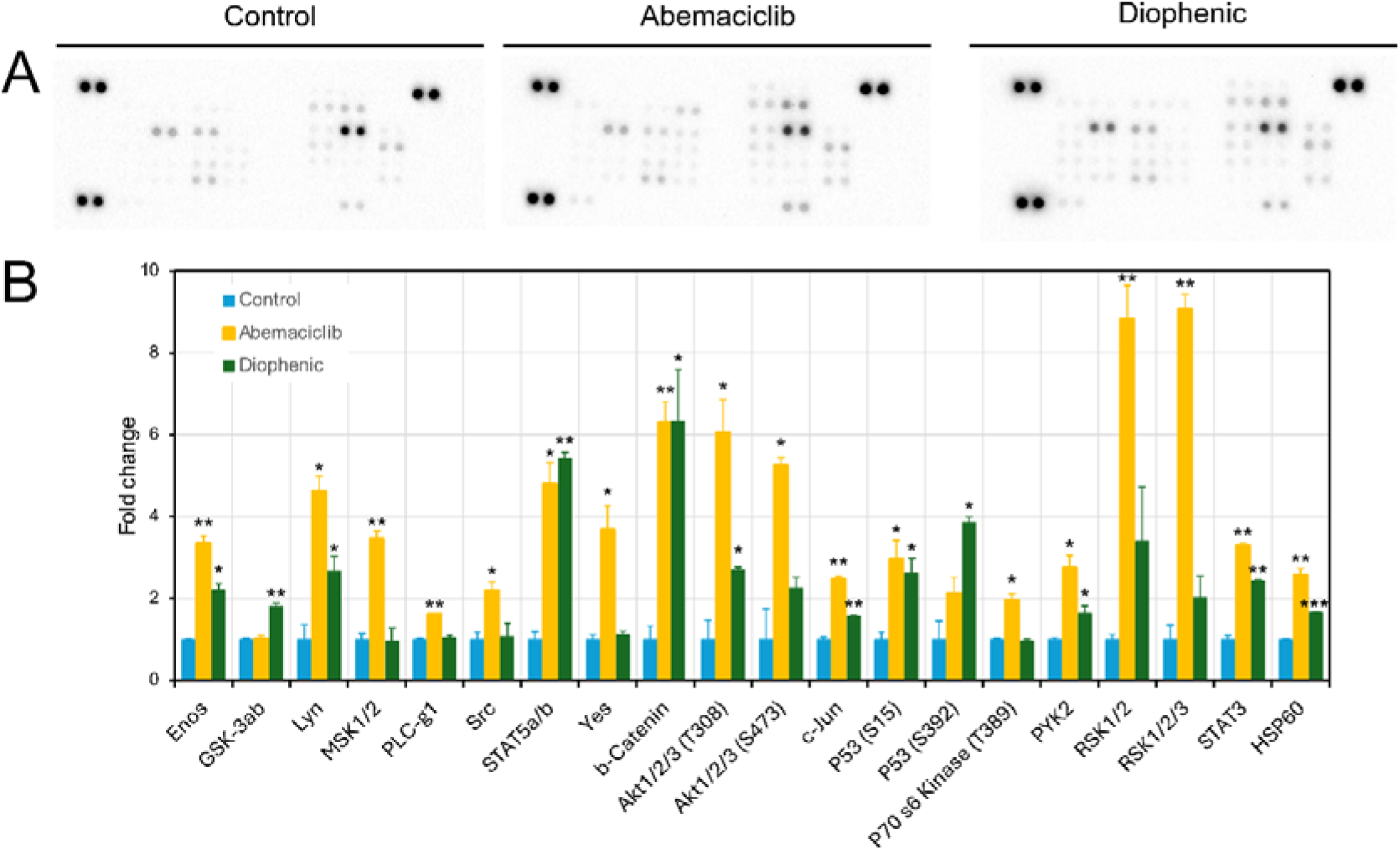
Effect of abemaciclib and diophenic on protein kinase activation in GB-42 cells. (**A**) Representative results from a human phospho-kinase array performed under control conditions or after 3 h of treatment with 25 µM abemaciclib or diophenic. Each condition includes the two membranes provided in the kit. (**B**) Quantification of fold-change relative to untreated control (n = 3). * *p* ≤ 0.05, ** *p* ≤ 0.01, *** *p* ≤ 0.001.

Many of the kinases selectively activated by abemaciclib have been described to be involved in stress response (i.e. MSK1/2, P70S6K, RSK1/2/3) and migration-associated signaling pathways (i.e. cJun, PYK2, PLC-γ1, eNOS, Src, Akt), which could explain the differential outcomes observed in the migration assay (**Fig. 6A**). These results suggest that abemaciclib may exert broader off-target activity beyond CDK4 inhibition. In contrast, diophenic modulates a more restricted subset of kinases, consistent with its design to interfere specifically with the CDK4-Cyclin D1 interaction rather than with the highly conserved ATP-binding domain.

Together, these findings indicate that diophenic acts as a more selective CDK4 inhibitor, displaying fewer off-target effects, a property that could underlie the distinct biological responses observed in previous experiments.

## 4. Discussion

The aggressive nature of GBM, marked by a median survival of only 12 months [1], underscores the urgent need for novel therapeutic strategies. Consistent with previous reports, our earlier study identified CDK4 overexpression in GBM [4], providing a strong rationale for the development of kinase inhibitors as a potential therapeutic strategy [53–55].

The FDA- and EMA-approved CDK4/6 inhibitors (i.e., palbociclib, ribociclib, and abemaciclib) have demonstrated efficacy in HR+/HER2− breast cancer. However, their mechanism of action, which relies on blocking the ATP-binding pocket, presents significant limitations [15,25–27]. This catalytic site is highly conserved across the kinase superfamily [47], resulting in limited specificity. Such non-selectivity is believed to contribute to common clinical toxicities, including neutropenia, nausea, and diarrhea [49,50]. While CDK4 has emerged as a key driver of GBM progression [4,53–55], the clinical utility of first-generation CDK4/6 inhibitors has been constrained by insufficient selectivity and toxicities associated with ATP-competitive inhibition [15,47,49]. Using an integrated computational and experimental approach, we identified diophenic, a novel PPI inhibitor that selectively disrupts the CDK4-Cyclin D1 complex while sparing the closely related CDK6-Cyclin D1 complex, thereby addressing a key limitation of conventional catalytic inhibitors.

We chose to investigate CDK4 as a potential therapeutic target in GBM because of its well-established role in cell-cycle regulation through RB phosphorylation and the subsequent release of E2F from the RB complex, a critical step in the G1-to-S phase transition that is considered the canonical pathway of CDK4 activity. Additional mechanisms include CDK4/6-mediated phosphorylation of the transcription factor FOXM1, whose degradation in the absence of phosphorylation increases ROS levels, a proposed priming mechanism for senescence [25]. Interestingly, we found that diophenic induces pronounced caspase-dependent apoptosis in GBM cells (**Fig. 5**), rather than the predominantly cytostatic and senescence-associated response typically elicited by ATP-competitive CDK4/6 inhibitors [20,21]. Commercial CDK4/6 inhibitors generally promote stable cell-cycle arrest through mechanisms involving the p16 INK4A/RB axis or FOXM1 degradation [25,29,56]. In contrast, the pro-apoptotic effects of diophenic suggest that disrupting the physical interaction between CDK4 and Cyclin D1 has distinct biological consequences. This shift from cytostasis to cytotoxicity may be particularly advantageous in aggressive tumors such as GBM, where effective tumor cell elimination is critical for reducing recurrence. This distinct outcome may reflect differences in the extent, kinetics, or cellular consequences of disrupting the CDK4-Cyclin D1 interface compared with ATP-competitive inhibition of CDK4/6, and warrants further mechanistic investigation.

It has been reported that breast cancer cells treated with CDK4/6 inhibitors secrete the chemokines CCL5 and CXCL10, thereby facilitating T-cell infiltration [12]. The CDK4-Cyclin D1 complex also plays an important role in cellular dynamics, including migration, invasion, and metastasis, across multiple cancer models [9]. This is partly achieved through the interaction of CDK4-Cyclin D1 with, or phosphorylation of, proteins involved in cytoskeletal regulation [57] and epithelial-to-mesenchymal transition (EMT) [58,59]. Our functional studies and phosphokinase profiling further highlight the enhanced functional selectivity of diophenic compared with ATP-competitive inhibitors. Notably, diophenic markedly reduced GBM cell invasiveness while exerting limited effects on cell migration in wound-healing assays (**Fig. 6**). This phenotypic dissociation was supported by phosphokinase array analysis (**Fig. 7**), which showed that several migration-associated signaling nodes, including Src, PLC-γ1, PYK2, and Akt, were modulated by abemaciclib but remained largely unaffected by diophenic. Given that CDK4/6-dependent cellular dynamics involve extensive phosphorylation of cytoskeletal and EMT-related targets [57–61], these findings suggest that ATP-competitive inhibitors perturb broader non-canonical signaling networks, whereas disruption of the CDK4-Cyclin D1 interface preserves essential basal signaling pathways while selectively impairing invasive tumor behavior.

The restricted signaling footprint of diophenic is also consistent with a favorable selectivity profile. By sparing the CDK6-Cyclin D1 complex, diophenic may offer a broader therapeutic window than currently approved CDK4/6 inhibitors such as palbociclib and abemaciclib [49,50]. Furthermore, given the established roles of CDK4 in metabolic reprogramming, mitochondrial regulation, and immune modulation [16,23,24,62,63], targeting the CDK4-Cyclin D1 interface may provide a valuable tool for dissecting canonical and non-canonical CDK4 functions while minimizing the off-target effects associated with global kinase inhibition.

In this context, the disruption the CDK4-Cyclin D1 interaction offers a promising solution to overcome the limitations of ATP-competitive inhibitors [13,14]. Since Cyclin D1 is the essential partner that determines CDK4 localization and oncogenic function, disrupting this PPI could provide enhanced functional selectivity [13,14]. Similar strategies have been proposed for other enzymes like the development of mTOR inhibitors directed against an allosteric regulatory site rather than the catalytic domain performed by our group [30,51].

Our findings suggest that the apoptotic response elicited by diophenic may reflect a specific dependency of GBM cells on the integrity of the CDK4-Cyclin D1 complex, rather than on CDK4 catalytic activity alone. This observation may help explain the distinct cellular outcome observed following interface disruption compared with ATP-competitive inhibition, which predominantly induces cytostatic responses. Although the precise molecular mechanisms remain to be elucidated, our results indicate that the structural and signaling functions associated with the CDK4-Cyclin D1 complex are critical for GBM cell survival. These findings further support the targeting of PPIs as a mechanistically distinct strategy capable of uncovering therapeutic vulnerabilities that may not be accessible through conventional kinase inhibition.

## 5. Conclusions

This study identifies the CDK4-Cyclin D1 protein-protein interface as a therapeutically actionable target in GBM, offering a potential alternative to catalytic ATP-competitive inhibitors. Through an integrated computational and experimental workflow, we identified diophenic, a novel candidate that selectively disrupts the CDK4-Cyclin D1 complex while sparing CDK6. Notably, diophenic bypasses the predominantly cytostatic response associated with ATP-competitive inhibitors, instead inducing robust caspase-dependent apoptosis and suppressing GBM cell invasiveness. Furthermore, phosphokinase profiling suggests that targeting this interface results in a more restricted perturbation of cellular signaling pathways, potentially contributing to improved functional selectivity. Collectively, these findings provide proof of concept for PPI-based CDK4 inhibition, supporting further optimization and *in vivo* validation of this strategy as a potential therapeutic avenue in GBM and other cancers.

## CRediT authorship contribution statement

**María Fuentes-Baile**: Methodology, Software, Validation, Investigation, Data curation, Writing—original draft preparation, Visualization. **José Antonio Encinar**: Molecular modeling, Virtual screening, Data curation, Writing—review and editing, Funding acquisition. **Salomé Araujo-Abad**: Methodology, Validation, Investigation. **Elizabeth Perez-Valenciano**: Methodology, Validation, Investigation. **Pilar García-Morales**: Methodology, Validation, Investigation. **Laura Fuertes-García**: Methodology, Validation, Investigation. **Camino de Juan Romero**: Conceptualization, Formal analysis, Resources, Writing—review and editing, Funding acquisition. **Miguel Saceda**: Conceptualization, Methodology, Formal analysis, Resources, Writing—original draft preparation, Funding acquisition.

## Data Availability

The datasets used and/or analyzed during the current study are available from the corresponding author on reasonable request.

## Ethics approval

Not applicable.

## Consent for Publication

Not applicable.

## Appendix A. Supplementary data

Supplementary data to this article can be found in Supplementary-file.pdf.

## Abbreviations

Δ*G*: Gibbs free energy variation
*ADMET*: Absorption, distribution, metabolism, excretion, and toxicity
*BSA*: Bovine serum albumin
*CCND1*: Cyclin D1
*CDK4*: Cyclin dependent kinase 4
*CDK6*: Cyclin dependent kinase 6
*COAD*: Colorectal carcinoma
*EMA*: European Medicines Agency
*EMT*: Epithelial-mesenchymal transition
*FBS*: Fetal bovine serum
*FDA*: Food and Drug Administration
*GBM*: Glioblastoma
*HER2-*: Human epidermal growth factor receptor 2-negative
*HGUE*: Hospital General Universitario de Elche
*HR+*: Hormone receptor-positive
*IDH1/2*: Isocitrate dehydrogenase ½
*IF*: Immunofluorescence
*IMIM*: Instituto Municipal de Investigaciones Médicas
*MD*: Molecular dynamics
*MGMT*: O6-methylguanine-DNA methyltransferase
*MM/PBSA*: Molecular mechanics Poisson–Boltzmann surface area
*MTT*: Methylthiazolyldiphenyl-tetrazolium bromide
*MW*: Molecular weight
*PAAD*: Pancreatic adenocarcinoma
*PBS*: Phosphate buffered saline
*PFA*: Paraformaldehyde
*PLA*: Proximity ligation assay
*PPI*: Protein–protein interaction
*RB*: Retinoblastoma
*RMSD*: Root-mean-square deviation
*SASP*: Senescence-associated secretory phenotype
*SD*: Standard deviation
*TME*: Tumor microenvironment
*TPSA*: Topological polar surface area

## Acknowledgements

We are grateful to the Centro de Computación Científica (CCC-UMH) for providing access to the Castleblack computing cluster (https://ccc.umh.es/). Diophenic and compounds 1 and 2 have been submitted for international patent protection (PCT/ES2025/070406; WO2026008907 A1 priority date: July 2, 2025) for their use as PPI inhibitors of CDK4/cyclin D and for the treatment of cancers characterized by CDK4/cyclin D overexpression.

---

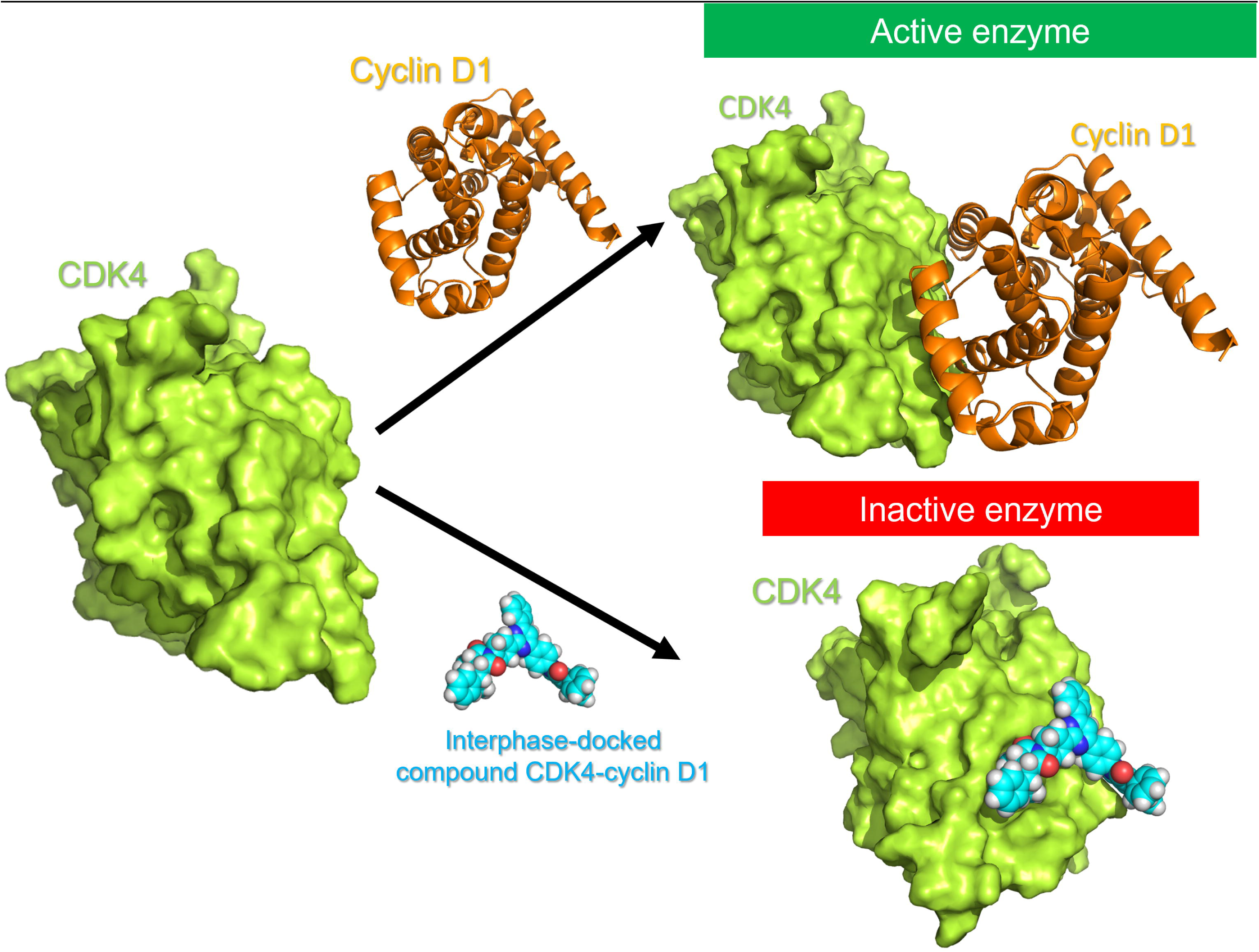

