## Supplementary data for "Targeting the CDK4-Cyclin D Complex: A New Generation of Selective Kinase Inhibitors for Cancer Therapy"

**Fig. S1. CDK4 expression levels in GBM and non-tumoral samples**. Data were obtained from RNA-seq results generated by our group (**A**), GlioVis (**B**), and UALCAN (**C**).

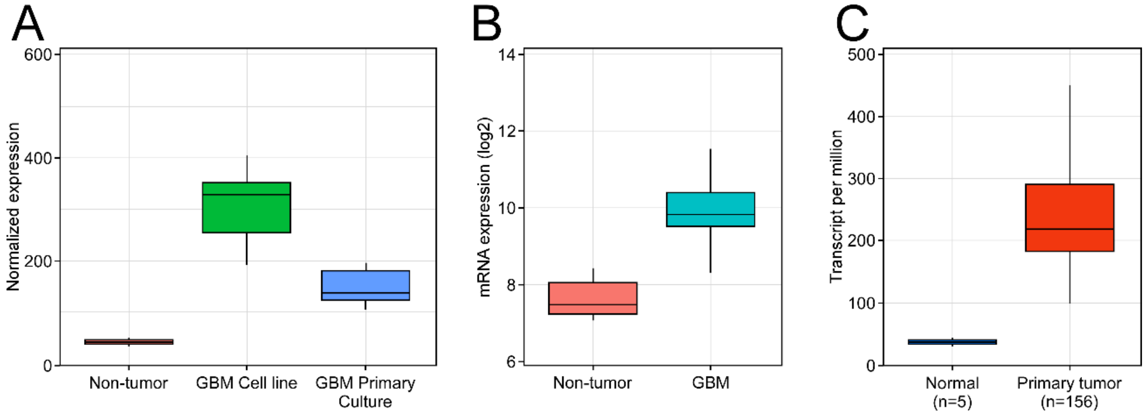


**Fig. S2. Dose-response curves for diophenic in GBM cell lines.** Cell proliferation, expressed as a percentage relative to the untreated control, was assessed after treatment with diophenic at concentrations ranging from 0.1 to 25 µM. Data are presented as mean ± SD (n ≥ 6) with the corresponding $IC_{50}$ values. Dose–response curves were fitted using a three-parameter logistic model [Y=100/(1+10^((X-LogIC50)), X: log of inhibitor concentration, and Y: normalized response, from 100 down to 0], and the legend indicates the corresponding $IC_{50}$ values.

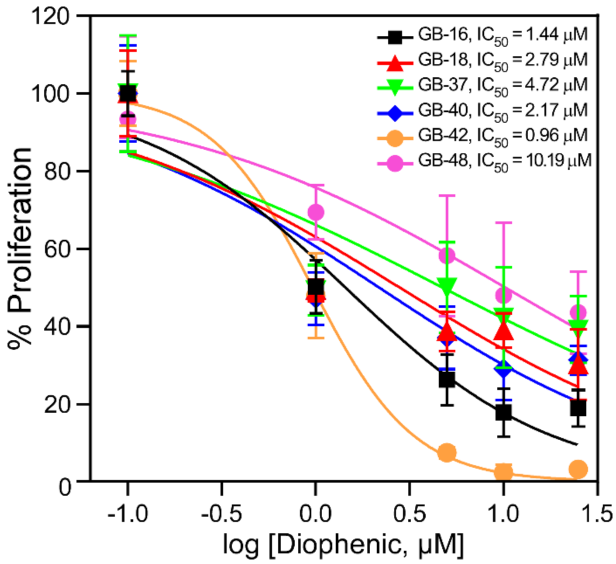

**Fig. S3. Immunofluorescence of CDK4 and Cyclin D1**. (**A**) CDK4 (red), Cyclin D1 (green), and DAPI (blue) in cancer cell lines. (**B**) Cytogram and correlation analysis of CDK4 and Cyclin D1 colocalization. The Pearson's correlation coefficient (PRV) and Manders' coefficients (M1 and M2) are shown, representing the fraction of Cyclin D1 (green) overlapping with CDK4 (red). PRV and Manders' coefficients (M1 and M2) were calculated using the JACoP plug-in in ImageJ. Scale bar: 10 µm.

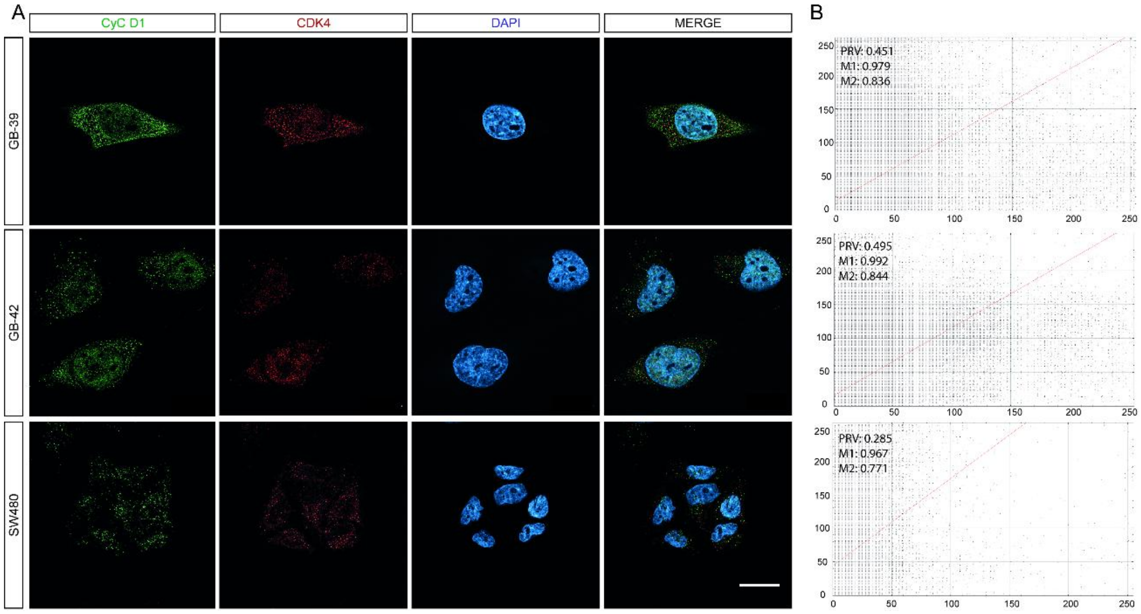

**Fig. S4. Effect of diophenic on the interaction between CDK6 and cyclin D1.** (**A**) Immunofluorescence of CDK6 and Cyclin D1 in GB-42 cells. Cyclin D1 (red), CDK6 (green), and DAPI (blue) are shown. (**B, C**) Proximity ligation assay (PLA) performed in GB-42 cells treated with 10 µM diophenic for 6 h. The scale bar represents 10 µm. Data are presented as mean ± SD (n ≥ 3).

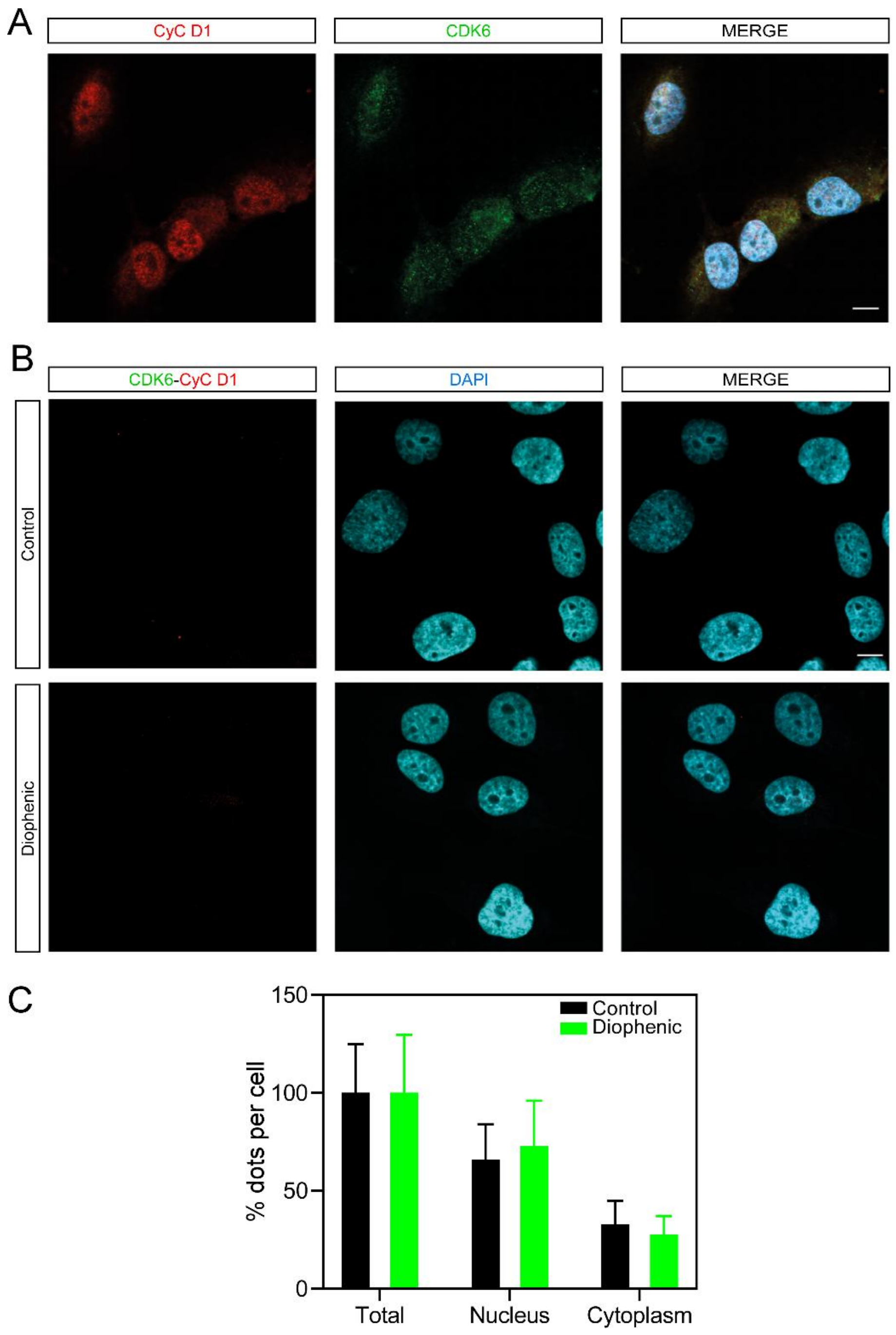

**Fig. S5. Effect of diophenic on the activity of E2F, Rb, and Myc.** GB-42 cells were transfected with pRb-TA-Luc, pE2F-TA-Luc, pMyc-TA-Luc, or the negative control vector pTA-Luc and subsequently treated with diophenic (10 and 25 µM) for 24 h. Luciferase activity was measured with the ONE-Glo EX Luciferase Assay system. Data are presented as mean ± SD of the relative luciferase activity normalized to the untreated control (n ≥ 3). ** $p \leq 0.01$, *** $p \leq 0.001$.

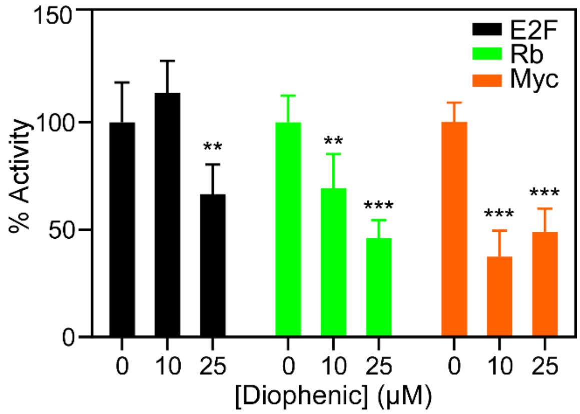


**Fig. S6. Effect of CDK4 silencing.** GB-16, GB-39, and GB-42 cells were transfected with a non-specific (NS) siRNA or an siRNA targeting CDK4. (**A**) CDK4 expression and (**B**) cell cycle distribution were analyzed. Data are presented as mean ± SD of CDK4 expression relative to the NS control and of the change in the percentage of cells in each cell cycle phase relative to the NS control (n ≥ 3). * $p \leq 0.05$, ** $p \leq 0.01$, *** $p \leq 0.001$.

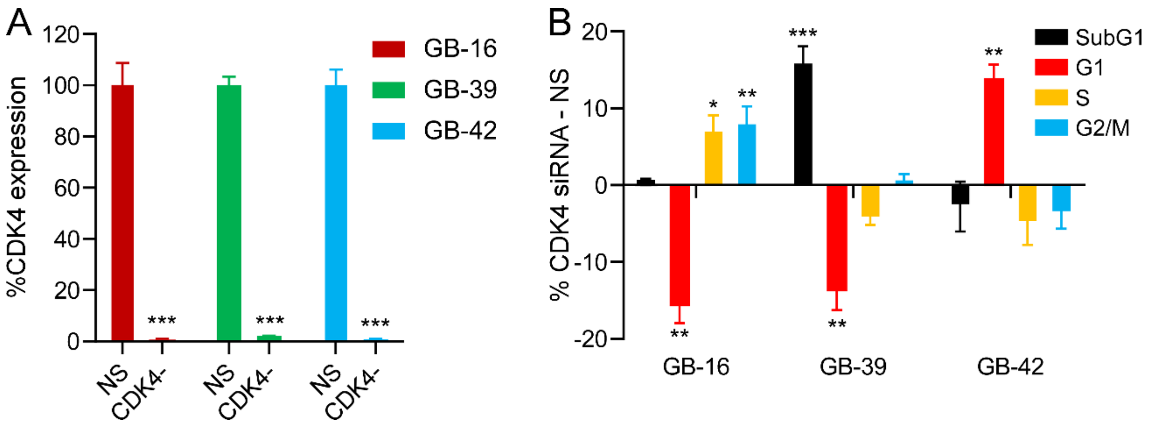

**Fig. S7. Migration assay with CDK4 inhibitors on tumor cell lines.** Representative images of the wound healing assay are shown for SW480**,** GB-39, and GB-42 cells treated with 25 µM diophenic, abemaciclib, or palbociclib for 48 h and labeled with Hoechst. Images were taken at baseline, 24, and 48 h. Scale bar: 1000 µm.

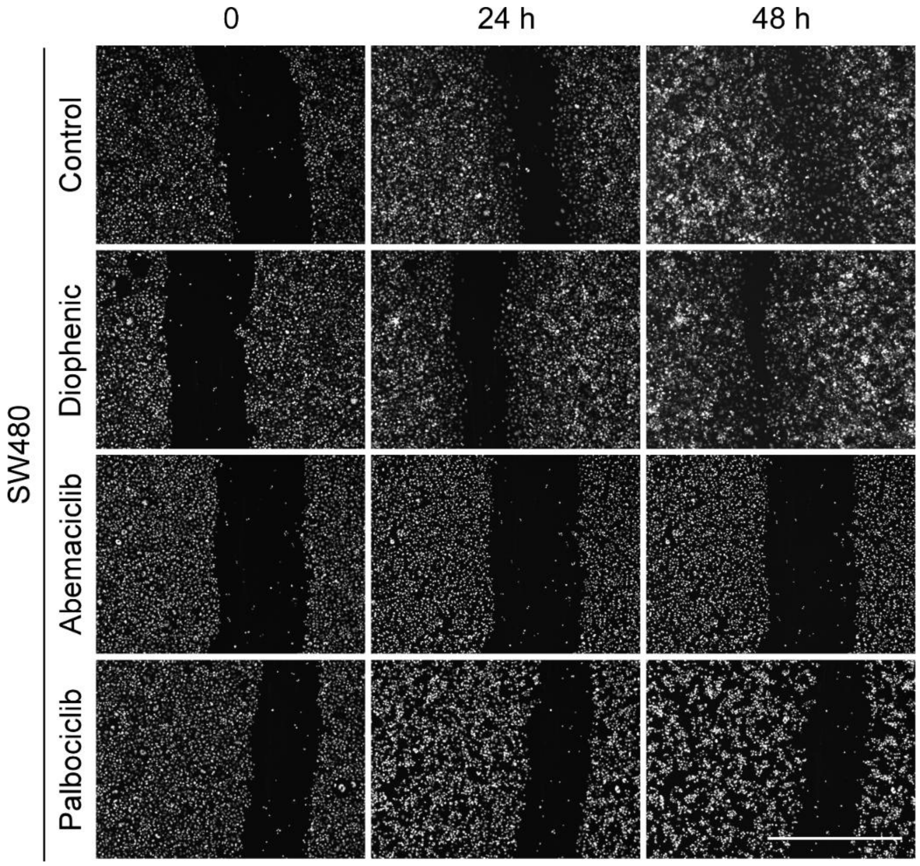

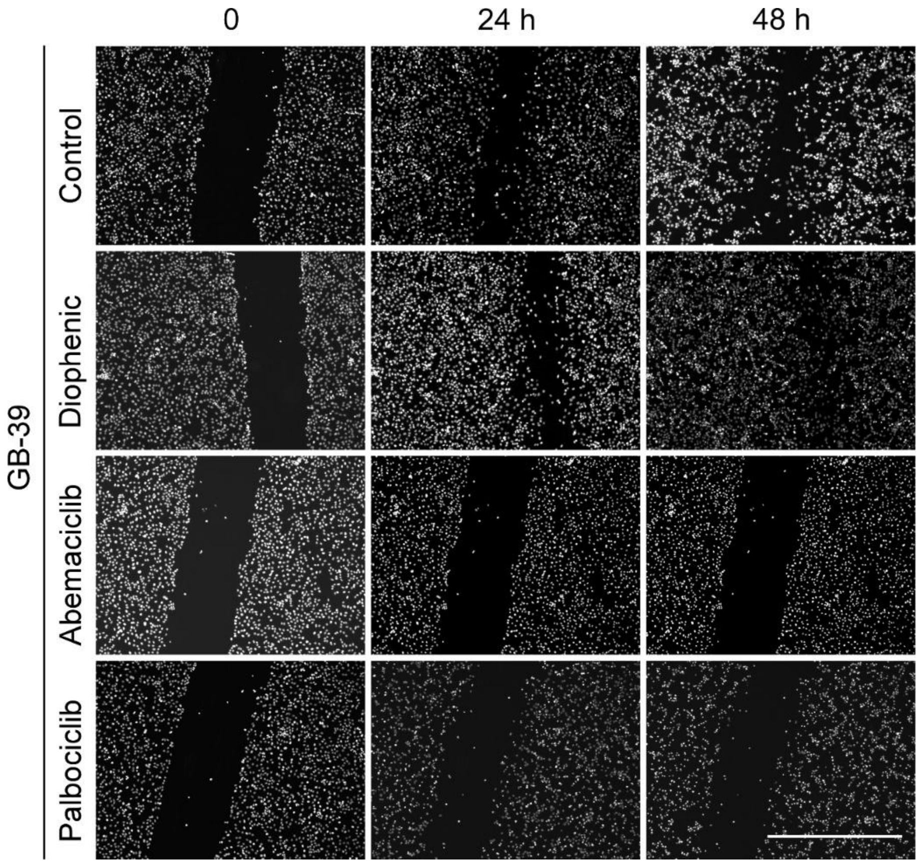
0
24 h
48 h
GB-39
Control
Diophenic
Abemaciclib
Palbociclib

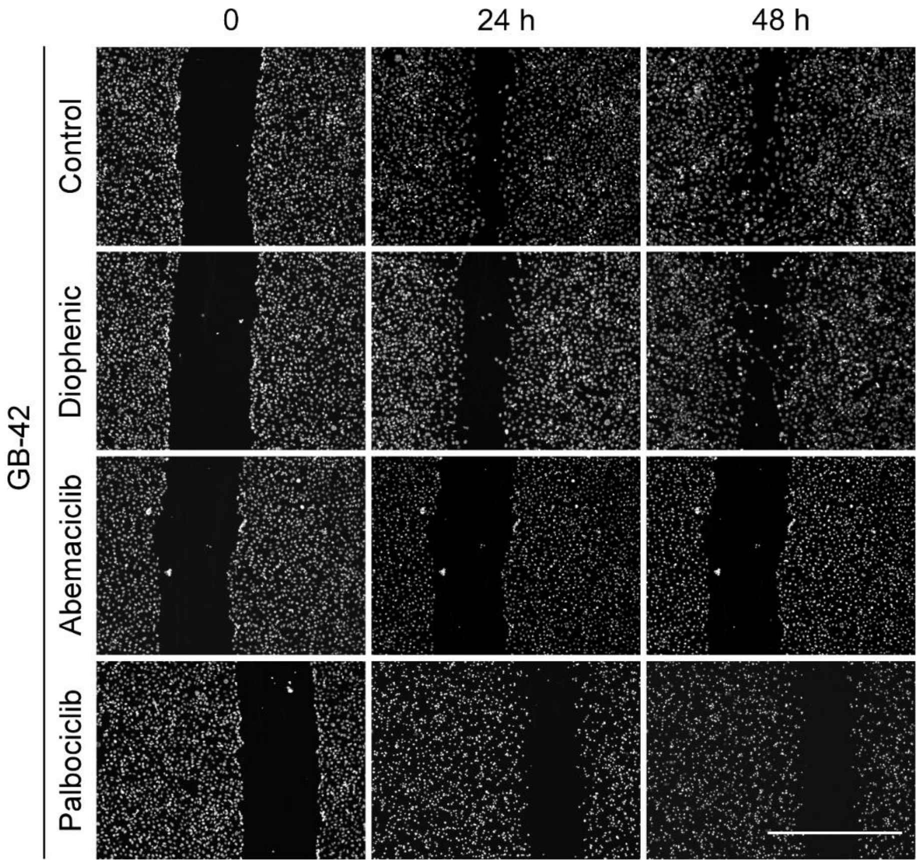
0
24 h
48 h
GB-42
Control
Diophenic
Abemaciclib
Palbociclib

**Table S1. List of candidate compounds selected as non-competitive CDK4 inhibitors**. The first column contains the identification number of each compound (used for reference throughout the manuscript). Columns 2 and 3 show the solvation binding energy values (MM/PBSA, kcal/mol) at the CDK4 interface throughout the entire MD simulation and during the last 50 ns, respectively. Column 4 presents the average ± standard deviation (SD) of the binding energy values calculated with Vina for each snapshot obtained during the MD simulation. Column 5 reports the maximum RMSD value observed in the trajectory of each compound during the MD simulation, representing the maximum displacement from its initial position. Column 6 provides the corresponding MolPort database name for each compound. The remaining columns include various descriptors and parameters, as indicated in their respective column headers.

| Number | MM\|PBSA 100 ns, Kcal/mol | MM\|PBSA last 50 ns, Kcal/mol | ∆G, Kcal/mol | Max RMSD, Å | Molport ID | Total Surface Area, Å$^2$ | TPSA | Rotatable Bonds | Aromatic Rings | cLogS | MW | cLogP | H-Acceptors | H-Donors | Ro5 violations |
|---|---|---|---|---|---|---|---|---|---|---|---|---|---|---|---|
| 1 | -43.191 | -42.358 | -7.882±0.7181 | 9 | *MolPort-001-024-020* | 386.34 | 74.76 | 3 | 6 | -10.622 | 544.565 | 7.366 | 6 | 0 | 2 |
| 2 | -46.191 | -46.782 | -8.05±0.6726 | 6 | *MolPort-002-134-975* | 462.78 | 95.58 | 7 | 6 | -10.666 | 613.671 | 8.1022 | 7 | 2 | 2 |
| 3 | -44.016 | -43.312 | -6.876±0.4657 | 4 | MolPort-002-754-815 | 400.67 | 98.01 | 5 | 4 | -7.024 | 607.467 | 3.9295 | 9 | 0 | 1 |
| 4 | -35.746 | -36.952 | -6.929±0.8429 | 4 | MolPort-001-014-219 | 371.65 | 66.48 | 3 | 5 | -8.136 | 520.587 | 5.2819 | 5 | 1 | 2 |
| 5 | -33.995 | -35.282 | -5.299±0.6626 | 9 | MolPort-001-547-847 | 513.84 | 83.98 | 7 | 6 | -10.462 | 660.859 | 9.2034 | 6 | 2 | 2 |
| 6 | -53.265 | -59.868 | -9.364±0.6676 | 4 | MolPort-001-548-163 | 479.7 | 66.4 | 4 | 6 | -8.894 | 618.779 | 8.7938 | 6 | 0 | 2 |
| 7 | -44.216 | -53.171 | -8.586±0.92 | 8 | MolPort-002-701-764 | 525.46 | 140.1 | 6 | 6 | -8.595 | 746.737 | 4.1255 | 12 | 0 | 2 |
| 8 | -34.489 | -40.438 | -5.798±1.042 | 7 | MolPort-002-751-769 | 445.97 | 98.01 | 5 | 4 | -7.604 | 630.653 | 4.5902 | 9 | 0 | 1 |
| 9 | -27.106 | -39.737 | -6.757±1.606 | 5 | MolPort-000-705-652 | 336.05 | 87.12 | 1 | 5 | -8.958 | 493.477 | 5.3592 | 7 | 0 | 1 |
| 10 | -59.504 | -54.604 | -8.223±0.6824 | 8 | MolPort-000-755-295 | 463.83 | 97.68 | 6 | 4 | -9.198 | 631.134 | 6.2236 | 9 | 1 | 2 |
| 11 | -36.639 | -34.866 | -6.997±0.6364 | 6 | MolPort-000-773-992 | 420.65 | 79.98 | 5 | 4 | -8.651 | 561.676 | 6.8775 | 6 | 1 | 2 |
| 12 | -33.006 | -38.674 | -7.323±0.6659 | 9 | MolPort-001-014-052 | 404.09 | 91.83 | 4 | 6 | -10.636 | 572.575 | 6.8368 | 7 | 0 | 2 |
| 13 | -60.106 | -71.37 | -5.273±0.5237 | 8 | MolPort-001-015-732 | 622.9 | 100.54 | 7 | 10 | -17.405 | 850.932 | 12.605 | 8 | 0 | 2 |
| 14 | -45.979 | -45.167 | -8.462±1.004 | 9 | MolPort-001-018-470 | 436.76 | 74.76 | 3 | 5 | -9.375 | 626.71 | 6.2821 | 6 | 0 | 2 |
| 15 | -34.753 | -45.865 | -6.623±1.091 | 5 | MolPort-001-022-836 | 551.28 | 129.72 | 11 | 6 | -10.942 | 721.767 | 8.0722 | 9 | 2 | 2 |
| 16 | -39.627 | -41.226 | -7.791±0.6741 | 18 | MolPort-001-025-403 | 462.78 | 95.58 | 7 | 6 | -10.666 | 613.671 | 8.1022 | 7 | 2 | 2 |
| 17 | -32.659 | -41.321 | -9.483±0.8404 | 6 | *MolPort-001-025-593* | 527.38 | 170.34 | 7 | 6 | -10.202 | 751.709 | 6.4422 | 13 | 2 | 3 |
| 18 | -41.487 | -44.479 | -8.038±0.9195 | 9 | MolPort-001-025-605 | 412.46 | 95.58 | 5 | 6 | -9.706 | 561.596 | 7.1726 | 7 | 2 | 2 |
| 19 | -32.811 | -36.806 | -7.037±0.8948 | 8 | MolPort-001-524-468 | 433.57 | 74.76 | 4 | 6 | -14.126 | 608.652 | 7.6276 | 6 | 0 | 2 |

| | | | | | | | | | | | | | | | |
|---|---|---|---|---|---|---|---|---|---|---|---|---|---|---|---|
| 20 | -32.933 | -37.59 | -6.803±0.5536 | 10 | MolPort-001-918-170 | 336.8 | 85.69 | 8 | 3 | -3.697 | 432.479 | 2.1719 | 8 | 1 | 0 |
| 21 | -35.017 | -35.796 | -7.145±0.6789 | 7 | MolPort-002-321-991 | 611.51 | 132.96 | 8 | 8 | -17.15 | 846.897 | 9.9226 | 10 | 2 | 2 |
| 22 | -35.654 | -39.519 | -8.24±0.5696 | 7 | MolPort-002-323-732 | 480.7 | 74.76 | 4 | 8 | -14.314 | 670.722 | 10.22 | 6 | 0 | 2 |
| 23 | -34.04 | -35.59 | -7.813±0.6859 | 12 | MolPort-002-515-701 | 300.94 | 79.77 | 2 | 6 | -8.717 | 417.423 | 7.0031 | 6 | 2 | 1 |
| 24 | -42.311 | -54.41 | -6.779±1.546 | 11 | MolPort-002-524-206 | 349.58 | 48 | 5 | 4 | -6.968 | 461.516 | 5.4431 | 5 | 0 | 1 |
| 25 | -39.774 | -35.974 | -6.56±0.8176 | 8 | MolPort-002-550-620 | 367.58 | 95.81 | 4 | 2 | -7.904 | 509.652 | 5.5966 | 7 | 2 | 2 |
| 26 | -47.627 | -55.911 | -7.649±1.048 | 12 | MolPort-002-938-642 | 551.28 | 129.72 | 11 | 6 | -10.942 | 721.767 | 8.0722 | 9 | 2 | 2 |
| 27 | -34.365 | -40.245 | -6.81±0.8115 | 6 | MolPort-003-842-843 | 408.3 | 66.7 | 6 | 5 | -11.688 | 524.622 | 6.9478 | 6 | 1 | 2 |
| 28 | -47.869 | -48.467 | -8.043±0.5565 | 7 | MolPort-007-571-043 | 385.19 | 45.55 | 4 | 5 | -7.127 | 509.607 | 5.9836 | 5 | 0 | 2 |
| 29 | -28.052 | -32.422 | -7.718±0.6388 | 8 | MolPort-035-874-010 | 371.33 | 88.75 | 2 | 3 | -4.774 | 507.592 | 3.4442 | 8 | 2 | 1 |
